# Do DNA and cytometric measures agree on genome sizes?

**DOI:** 10.1101/2025.11.26.690794

**Authors:** Donald G. Gilbert

**Affiliations:** Biology Dept., Indiana University, Bloomington, IN, USA

## Abstract

Measurement of DNA contents of genomes is valuable for understanding genome biology, including assessments of genome assemblies, but it is not a trivial problem. Measuring contents of DNA shotgun reads is complicated by several factors: biological contents of genomes, laboratory methods, sequencing technology and computational processing. This compares and shares complications with cytometric measures of genome size and contents. There is an obvious discrepancy between cytometry and current long-read assemblies: assemblies average significantly below cytometric sizes.

Measures of population changes within a species will control some of these complications. This report examines five species population sets with published cytometric and DNA data sets: *Arabidopsis thaliana* and *A. arenosa*, Arctic plants of *Cochlearia* genus, clonal populations of a rotifer *Brachionus asplanchnoidis*, and *Zea mays* corn plants. Results of this are clear, if complicated: DNA and cytometry do measure the same genome sizes, when done carefully with controls or adjustments for errors.

Population changes in genome sizes are found by assembly-mapped measures of DNA, including environment or regional effects of latitude and altitude. Copy numbers of repeats, transposons and genes are changing. Kmer-based measures of DNA generally fail to match cytometry, miss population changes, and are opaque to understanding measurement errors. Oxford Nanopore technology produces the least biased DNA for measurement, with recent ONT.R10 Simplex data a match to cytometric sizes for corn, tomato plants, zebra fish and zebra finch bird. Assemblies of these species DNA average 12% below measured sizes, incomplete for duplicated content. New assembly of this ONT Simplex DNA reaches the size measured by cytometry and Gnodes, in 4 of 5 species. rRNA gene duplications measure one aspect of this discrepancy: the genome assemblies examined all are missing many rRNA genes. New experiments that measure both cytometry and DNA, controlling error factors, are warranted to clarify these results and suggest improvements for genome projects.

## Introduction

Cytometry and DNA sequencing offer two independent means to measure nuclear genomes. There is limited experimental verification that cytometry and DNA sequencing measure the same nuclear genome sizes, and often large discrepancies between them (e.g. Pflug et al 2020; Mgwatu et al 2020; Li et al 2025). Cytometry, measured from florescence of stained nuclear chromosomes, has well established protocols for genome measurements (Galbraith et al 1983; Bennett et al 2003; Leitch et al 2019; Pellicer et al 2021; Sliwinska et al 2022). DNA sequencing, used to assemble the pseudo-molecules representing nuclear genomes, measures size by counting randomly sequenced nucleotide bases divided by sequencing depth (Lander and Waterman, 1988). Methods for DNA measurement (Vuture et al 2017; Sun et al 2017; Pucker 2019; Pflug et al 2020) are now widely used but are not as extensively validated as cytometry.

Recent telomere-to-telomere genome assemblies of many plants and animals are smaller than published cytometry sizes, by an average 12%, with some much smaller (Gilbert 2023, 2024). A majority of assemblies are from 10% to over 50% below cytometric sizes, with the same average 12% discrepancy for five species examined in detail (human, maize, zebrafish, sorghum, and rice) as for 500 species. Similar results of this cytometry - assembly difference are reported by Hjelmen (2024). Bennett et al. (2003) examine discrepancies in early *Arabidopsis* genome assembly and cytometric measures, with insights that remain valid 20 years later, including uncertainty that this model plant has a completed genome assembly.

It is time to resolve this discrepancy with experiments. Claims of complete genome assemblies need validation in the face of contrary evidence, as do claims that cytometry is accurate or flawed in its measures of genome sizes. The focus of this paper is to determine, from published data, if cytometry and DNA samples measure the same genome size changes, across several populations of one or related species. The best evidence for agreement of DNA measures with other genome size measures are studies using the same bio-samples, controlling biology, laboratory effects as much as feasible. A few such studies have been undertaken, notably with *Arabidopsis thaliana* model (Long et al 2013) and *Zea mays* inbred plant lines (Hufford et al., 2021). These are limited and ambiguous; early *A.t.* DNA sequences were unreliable measures, biased in GC content.

The discrepancy of techniques for genome measurements has been attributed by some authors to an an over-estimate by cytometric methods (eg. Sun et al 2017), but the argument for this is weak: since k-mer DNA estimates match assembly sizes, they must be correct, without providing evidence of biases in cytometry. Under-estimates, as well as over-estimates, by cytometry are likely, if the main error is in size estimates of standards (eg. Temsch et al. 2021). For example, report on maize inbred lines (Hufford et al., 2021) uses cytometry under-estimated by 10-15%, compared to data reported in its cited references (Chia et al 2012, cited by Wang et al 2021 and Hufford et al 2021). In contrast, under-estimates of genome size in DNA measures and assemblies are common, and detailed elsewhere (Gilbert 2022-24, Mgwatyu et al. 2020; Li et al 2025). Prior results (Gnodes #2) shows some agreement of DNA and cytometric measures; this follow-up looks in detail at available evidence.

## Methods, briefly

Recent efforts by this author have focused on accurately measuring genome DNA contents with genome statistical software termed **Gnodes**, a Genome Depth Estimator (Gilbert 2022-2024). This includes the problematic area of duplicated and highly repetitive structure in genomes. Results to date can inform new and updated genome projects with details of assembly completeness. Measuring or estimating DNA coverage depth is the critical problem for genome size measurement, with both assembly-mapped (Gnodes; MGSE: Pucker 2019; Pflug et al 2020) and k-mer methods (GenomeScope: Vuture et al 2017; findGSE: Sun et al 2017). Cytometry GSE relies on analogous statistical methods of finding peaks in the distributions of florescence quantities.

The equation G = LN/C (see Table DC1b Abbreviations) is used for DNA-GSE, with a variant used by k-mer methods. For Gnodes, two D depth, or C coverage, estimators are suggested for the results presented here: (1) D.ucg from known unique sequences measured with unbiased DNA, (2) D.all is a best estimator of depth for biased DNA, as the UCG are a very small portion of genomes. DNA read bias in GC content or CDS versus repeats will cause a large shift in read depth for the major portion of genome, so that depth over all spans better estimates it. This bias can be estimated as the ratio Cau = C.all/C.ucg; bias is suspect if (Cau < 1), no bias is estimated if Cau >= ∼1, as duplicated genome parts will increase Cau > 1. Gnodes software now reports such apparent DNA sequence bias.

This report examines five species-population sets that have published cytometric and DNA data sets: *Arabidopsis thaliana* and *Ar. arenosa*, Arctic plants of *Cochlearia* genus, rotifer *Brachionus asplanchnoidis*, and *Zea mays* corn plants, are used for the primary evidence of this work. Table DC.1 lists the published Cym and DNA data sets used in this study, including references of DNA and Cym data. Supplement tables include sequence read archive IDs for DNA examined, and the GSE values measured.

**Table DC1.**
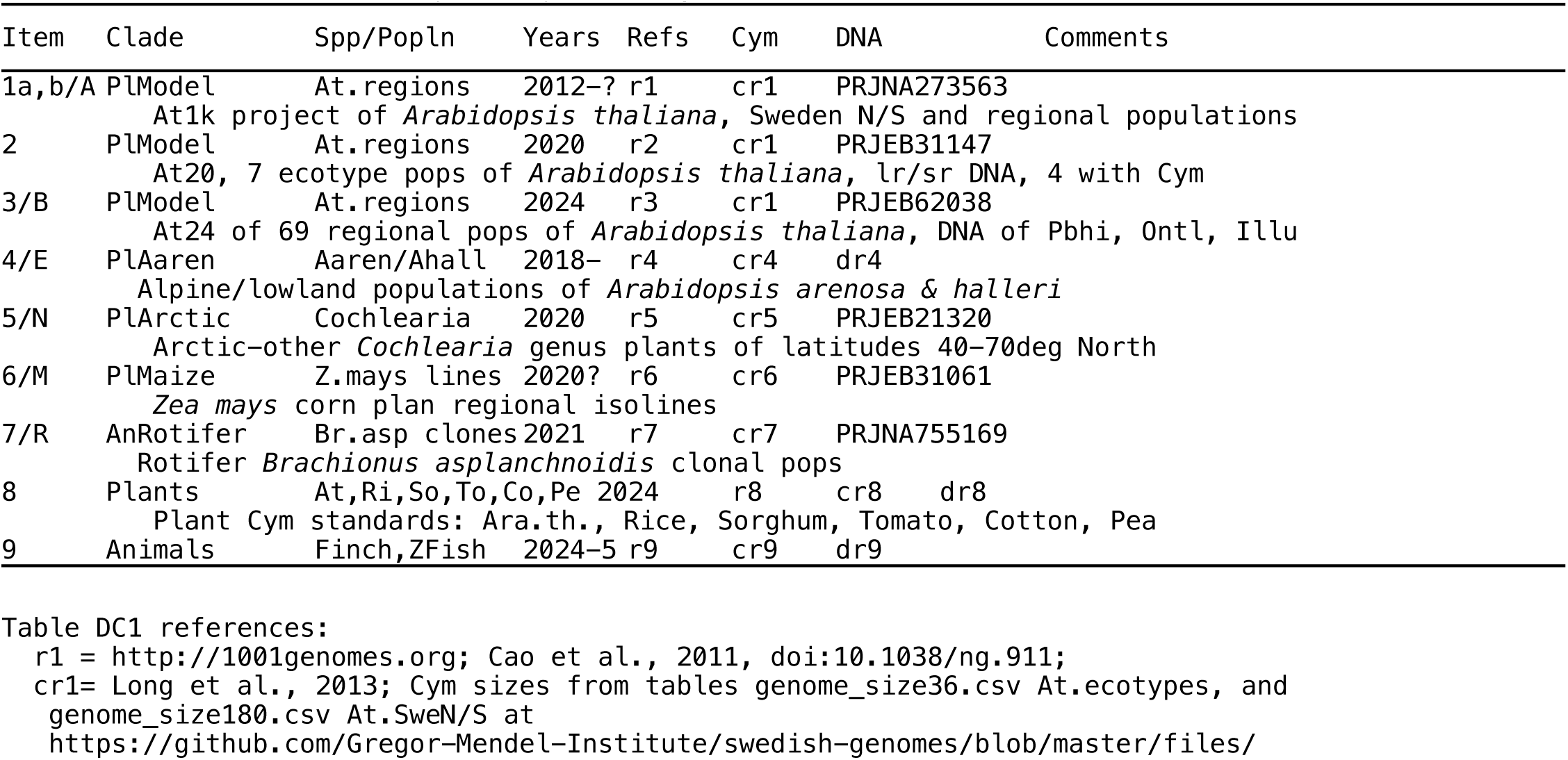
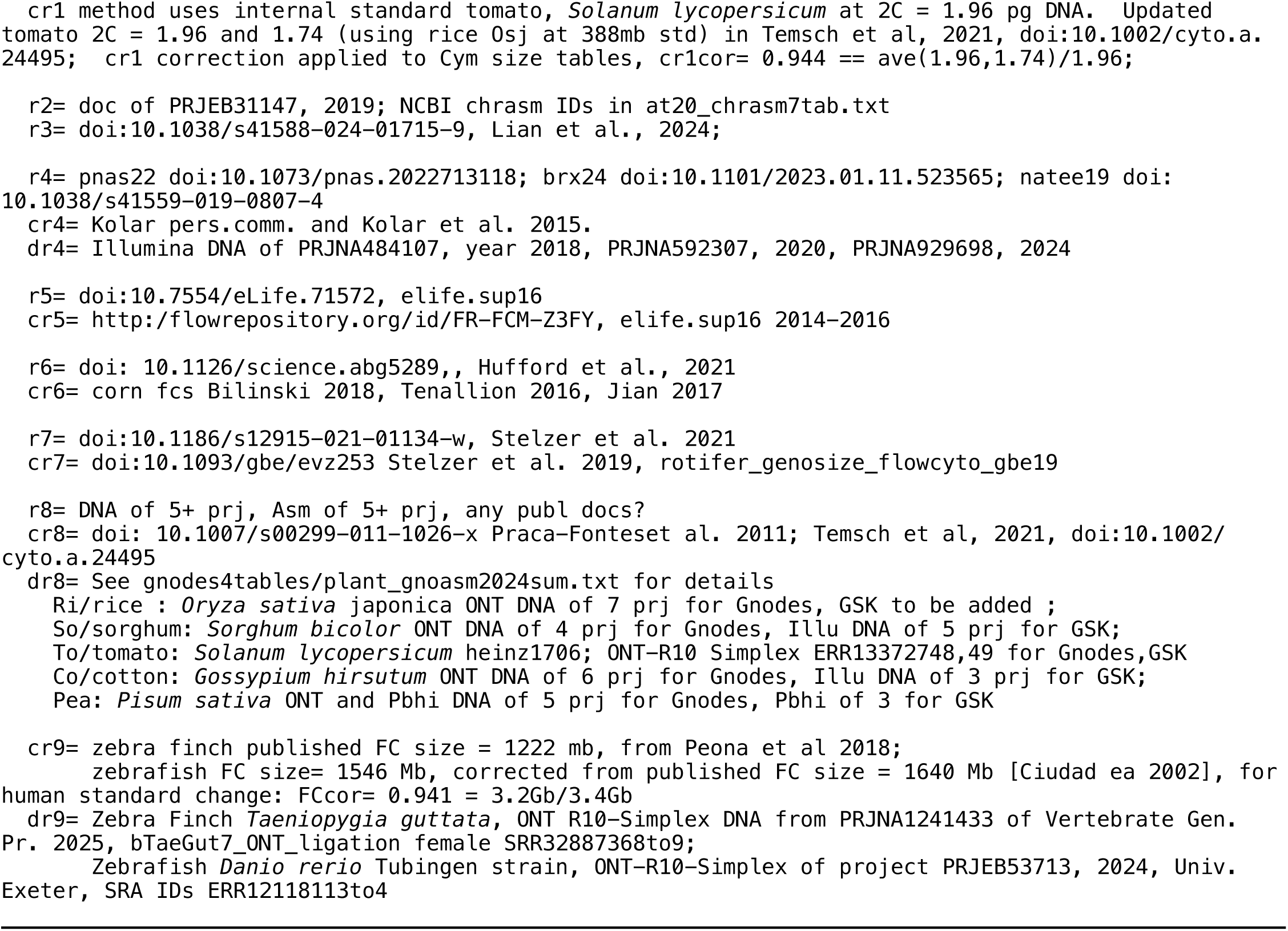
Data sets used in Cytometry x DNA genome size measures.

Genome assemblies published from the DNA read sets are used for measuring read coverage and copy numbers, with annotations of genome contents for transposons, coding genes, rRNA genes, satellite repeats, tandem repeats and other as produced by those projects or this project. Non-nuclear DNA has been removed: chloroplast, mitochondria and foreign contaminants identified by NCBI SRA reports (Katz et al 2021) and/or Kraken (Wood et al 2019).

**Table DC1b.**
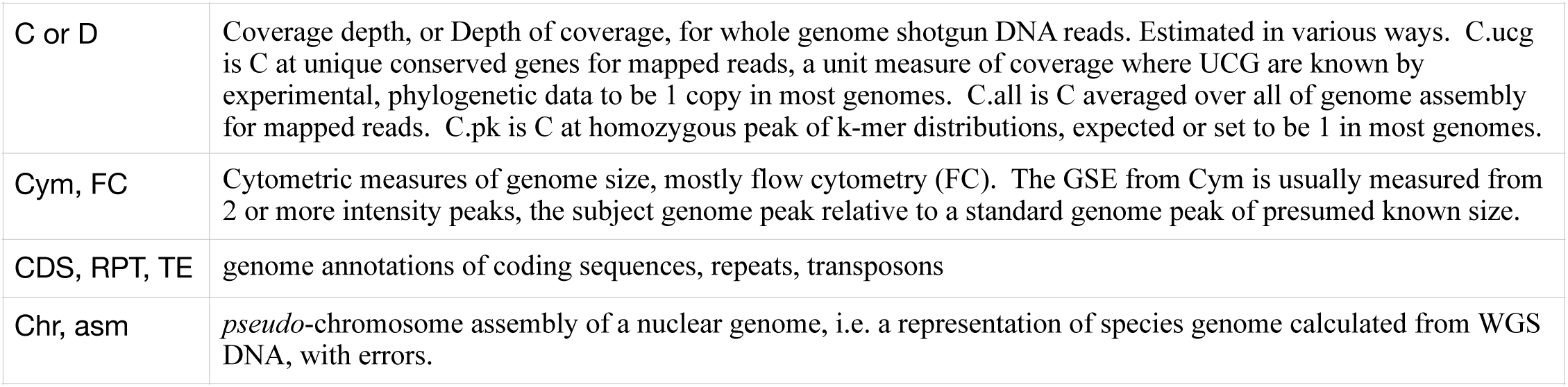
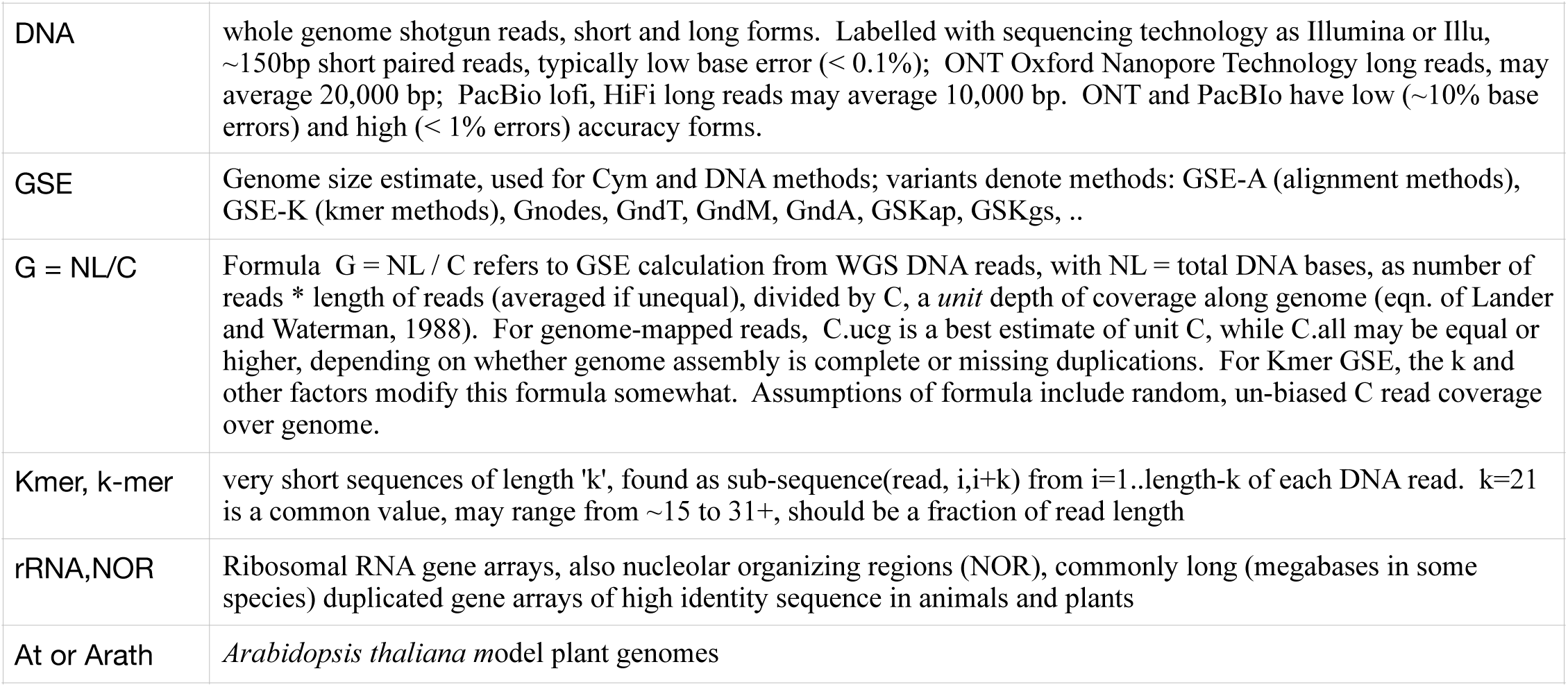
Abbreviations used here.

## Results in brief

The relation of genome sizes measured by DNA and cytometry is 1 to 1 over the large range of 150 to 4000 megabases for species examined, as shown in Figure DC1. The regression slope is 0.999 for log(DNA) x log(Cym), with an intercept of 1, indicating DNA and Cym measures are essentially the same values. This relation holds, but with more error, for the 5 population sets examined. There are small deviations: rice *O.s. japonica* has a 10% higher Cym (440mb) than DNA and 1:1 indicate (400mb). This relation over a large species range doesn’t address precision to detect smaller effects in genomes. The main question addressed here is whether population-level changes, on the order of 10%, are measurable and equivalent with DNA and Cym methods. If so, then assemblies of those DNA should take into account this completeness evidence.

**Figure DC1.**
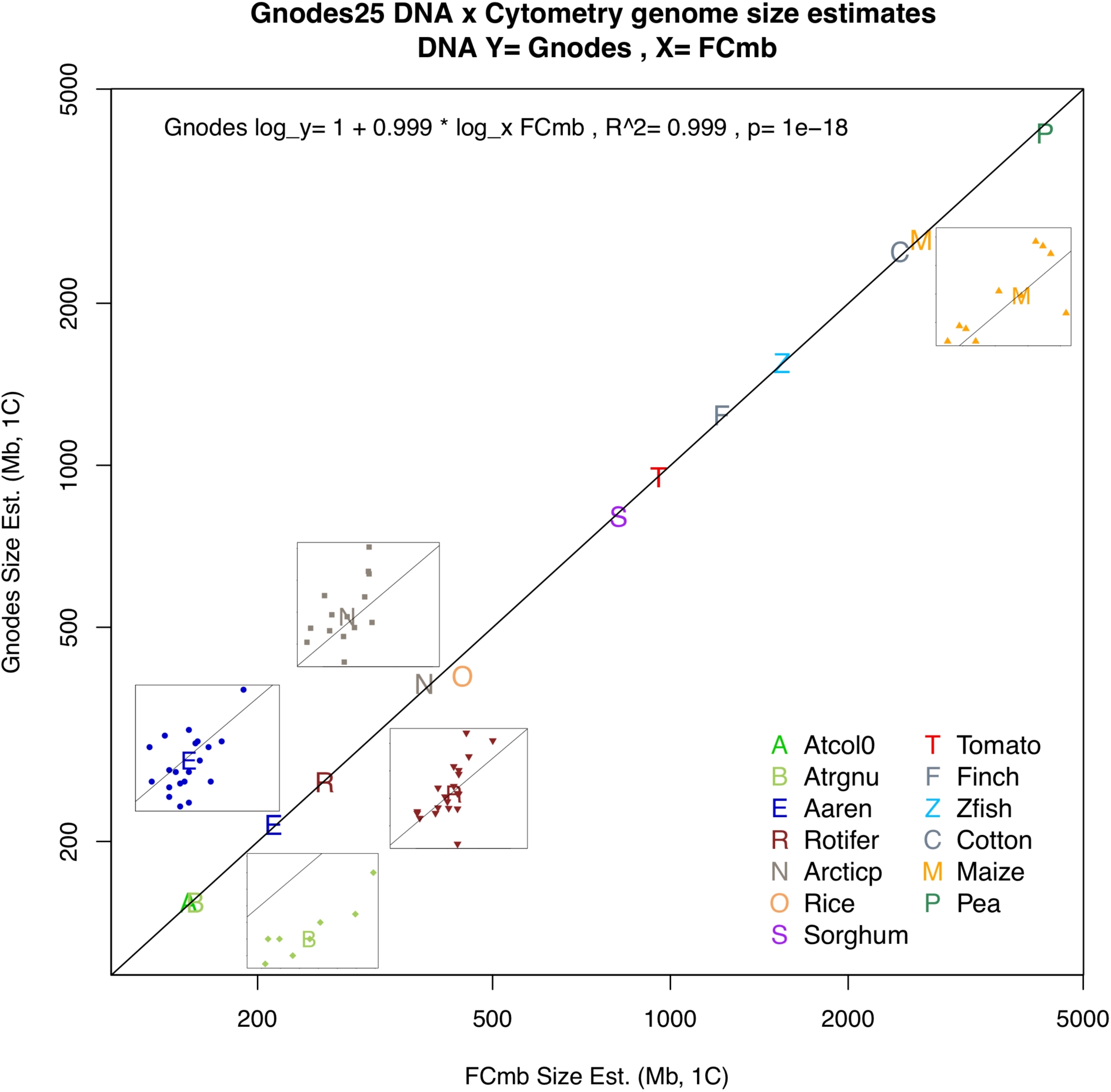
Regression of DNA x Cytometry genome sizes, measured for 5 species-populations along with 8 reference animals and plants, over the size range of 150 to 4000 megabases, on log XY-scale. Population variation is shown in sub-plots, not on same XY-scale, for lettered sets B,E,R,N,M of Table DC2. The relation is 1:1 (regression slope = 0.999 for logDNA x logCym). For population subsets, more error is seen, but all are statistically equal to 1:1 relations. Detail plots of these 5 follow in results detail section.

A general summary, in Table DC2, indicates that DNA measures equal Cym for all populations examined, while assembly sizes, where available for population sets, are mostly below Cym. Effects of latitude, altitude, regional populations, and laboratory methods are noted. Major contents of genomes that change across populations are also indicated: repeats, transposons, and gene coding sequences. Table DC3 itemizes changing genome contents related to regional effects, with reduced sizes for increasing altitude and latitude. Notably, repeats are changing for all species, CDS for 3 of 5, and large transposon changes in maize lines. Table DC4 indicates the amounts of regional effects for three species-population sets where these effects were found.

**Table DC2.** Result summary of Cytometry x DNA genome size measures. C=D, cytometry equals DNA genome measure; A=D assembly equals DNA measure; Effects: environmental, population or other factors with signif. genome size changes; Kmer: DNA measures by Kmer methods do not (No) match Cym; ContentDelta: major components of genome that change.

| Item | SppID | Npop/ind | C=D | A=D | Effects | Kmer | ContentDelta |
| --- | --- | --- | --- | --- | --- | --- | --- |
| 1a/A | At1kSwe | 42/42 | Yes~ | .. | N-S.Sweden | .. | RPT,TE |
| 1b/A | At1kRgn | 24/24 | Yes | .. | Lat,Lng,Alt | No | RPT,CDS |
| 2 | At20Rgn | 7/7 | Maybe | No | Regions | .. | .. |
| 3/B | At24Rgn | 65/103 | Yes | No | Latd,lab | No | RPT,CDS |
| 4/E | Aaren | 42/250 | Yes | No | Altitude,lab | No | RPT,CDS |
| 5/N | Arctic | 14/14 | Yes | .. | Latitude | Fail | RPT,TE |
| 6/M | Maize | 12/na | Yes | No | Pops | No | TE,RPT |
| 7/R | Rotifer | 21/21 | Yes | .. | Pops,lab | No | RPT,TE,CDS |
| 8 | Plants | 6/na | Yes | Mixed | Species | No | .. |

Notes: 1/A. srDNA:poor, strong bias; 2. lrDNA and srDNA, limited Cym; 3/B. lrDNA, srDNA, lab effect, older Cym limited to npop=8/65; 4/E. srDNA:bias, contam, lab effect, Cym limited to npop=21; 5/N. srDNA:bias,contam; 6/M. lrDNA:ok, srDNA:bias; 7/R. srDNA:bias, contam?, lab effect; 8. lrDNA only used, well studied;

### Major Contents and Regional Effects in Population Genome Sizes

The main effects in genome contents and regions for population genome size changes are noted in Tables DC3 (coding sequene, repeats and transposons), DC4 (latitude, longitude, altitude changes) and Figure DC3, showing regional clines in *Ar. thaliana* and *Ar. arenosa*.

**Table DC3.** Major contents effects of population genome size changes, from regression of DNA content measure on total size measure. Cym, DNA are total size measures (median values, as megabases of 1C/haploid) of cytometry, DNA by Gnodes; CDSann, RPTann, TEann are annotated contents of coding genes, repeats and transposons, ‘ns’ indicates non-significant effect, others are significant; Range.Mb has 10% to 90% quartiles of low, high total sizes; Npop is number of populations (Table 2).

| SppID | Cym | DNA | Npop | Range.Mb | Range Changing Contents |  |  | Notes |
| --- | --- | --- | --- | --- | --- | --- | --- | --- |
|  |  |  |  |  | CDS% | Repeats% | TEann% |  |
| B Atrgnu | 157 | 154 | 65 | 151,162 | 25 | 64 | 2.8ns | Latitude |
| E Aaren | 213 | 215 | 41 | 200,231 | 2.8 | 26 | 4.6ns | Altitude |
| R Rotifer | 260 | 258 | 19 | 211,315 | 4.1 | 57 | 6.9 |  |
| N Arcticp | 383 | 393 | 14 | 356,463 | 0.6ns | 8 | 7.3 | Latitude |
| M Maize | 2880 | 2794 | 12 | 2637,2965 | 3.7ns | 16 | 94 |  |

Notes: B. Good annotation, Latitude -cor Total,CDS,RPT; E. Altitude -cor Total,RPT +cor TE; N. Poor annotation, Latitude -cor Total,TE; M. Good annotation;

**Table DC4.** Regional effects (latitude, longitude, altitude) correlated with genome size measures: GTotal = Gnodes total size, Repeat, TEs, CDS = Gnodes major content sizes, GSKgs = Kmer-GenomeScope, GSKfi = Kmer-findGSE. Significant +/- effect correlation is given, and genome size ranges (Mb) are estimated from regression on effects.

| SppID | GTotal | CDS | Repeats | TEs | GSKgs | GSKfi | Effect |
| --- | --- | --- | --- | --- | --- | --- | --- |
| B Atrgn | -North<br>158-154 | -North<br>52-51 | -North<br>28-25 | ns<br>19-20 | -North<br>144-139 | -North<br>144-138 | Latitude |
| E Aaren | -Alti<br>226-206 | ns<br>39-39 | -Alti<br>38-23 | +Alti<br>21-30 | ns<br>182-178 | fail<br>.. | Altitude |
| N Arcti | -North<br>445-351 | -North<br>6-5 | -North<br>26-22 | -North<br>56-45 | fail<br>N0ann=323-257 | fail | Latitude |

Notes: B. Est. range for Latitude= 0-60 deg North; E. Est. range for Altitude= 0-2000 meters; N. Poor annot: N0ann contains CDS,RPT,TE; Est. range for Latitude= 40-70 deg North;

Figure DC3 shows the significant effects of latitude and altitude on population genome sizes, measured from DNA by Gnodes, for *Ara. thaliana* (DC3.B), and *Ara. arenosa* (DC3.E). For *Ara. arenosa*, there are 5 locales with nearby alpine and foothill populations, where the alpine ones are recently evolved from the foothills [cite https://doi:10.1073/pnas.2022713118]. The short lines show a cline of reduced genome sizes with altitude, for 4 of these pairs, in common with general trend.

### Quality of several DNA measures relative to Cytometry

Qualities of agreement for several DNA measures with Cym are summarized in Table DC5. Aspects of agreement are difference from Cym, correlation of DNA with Cym across populations of a species (ignoring average difference), and significance of a 1:1 relation of DNA measures with Cym. This summary finds the mapped DNA measure of Gnodes agrees with Cym measures, while the Kmer-based measures are not in agreement.

**Table DC5.**
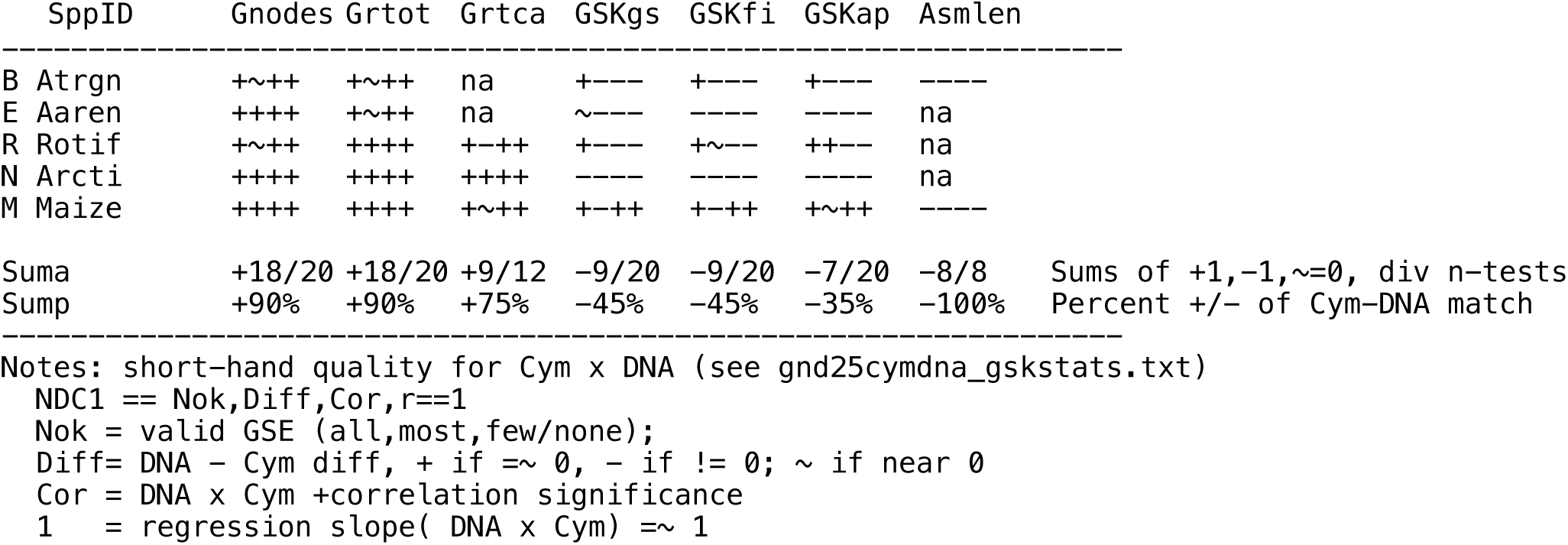
Quality of Cym x DNA for Gnodes and Kmer DNA measures. Quality string is NDC1 = Nok,Diff,Cor,Slope.r=1 with values of + pass, - fail, ∼ near-pass, na is missing, using statistical significance of Diff=t.test(Cym-DNA), Cor=cor.test(DNA∼Cym), Slope=regression(DNA∼Cym).slope == 1, Nok= Nvalid/Nsamples. DNA Kmer methods are GSKgs GenomeScope, GSKfi findGSE, GSKap basic kmer-peak where GSKap= sum(fk*nk)/ Cpeak; Gnodes measure Grtot is total DNA, and Grtca is bias-corrected.

Table DC6 examines only ONT R10 Simplex samples, due to the reported un-biased genome sampling of this method [Oxford Nanopore Technology, 2025; Lerminiaux et al 2023]. Two vertebrates are included here, the bird zebra finch, similar to chicken in size measures, and zebrafish. In this limited sample of ONT R10 Simplex DNA, Gnodes and Cym values are very close. Kmer GSE is possible with these data, the results are similar to short read DNA: GSKgs under-estimates Cym sizes by 5% to 13%, GSKfi has a broader range of under and over estimates. However, the simple GSKap formula [Lander & Waterman, 1988; Li & Waterman 2003] matches Gnodes and Cym more closely. The ONT R10 Simplex DNA method has read accuracy at or above 99% as well as the low bias in measuring genome content types of earlier ONT technology (eg. see Gnodes#2, esp. Figure F2 of rRNA and other repeat/duplicated contents). This is in contrast to ONT R10 Duplex (2-strand) result for Maize, produced from these same Simplex (1-strand) read sets, where rRNA and TE duplications are reduced or filtered out.

**Table DC6.**
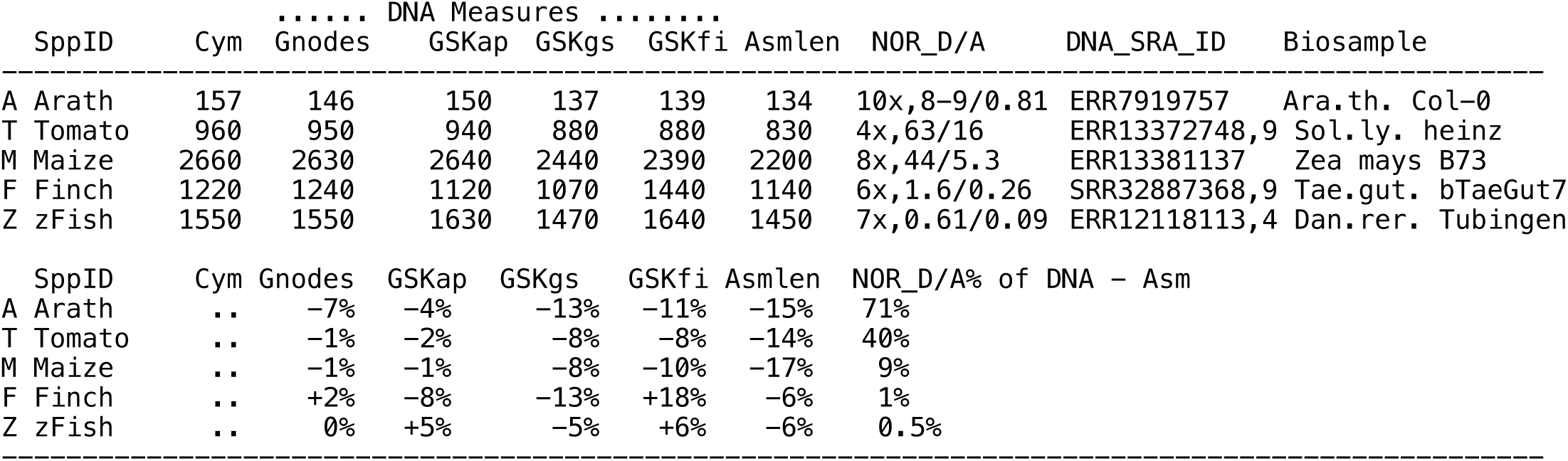
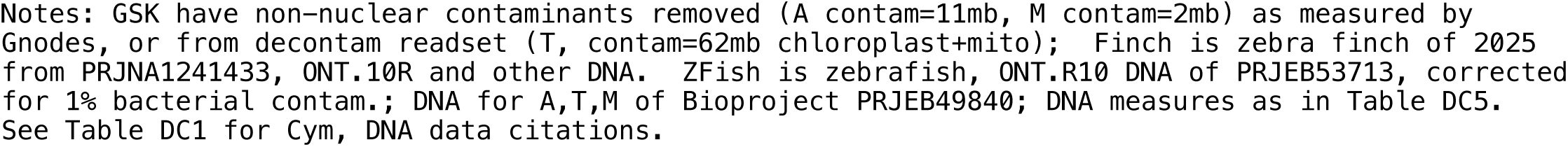
Genome sizes measured in DNA from high accuracy, low-bias method of ONT R10 Simplex, and standard cytometric size (Cym), as megabases (1C, haploid size), part A. Part B is percent deviation from Cym, but NOR_D/A% is percent of DNA - Asm that is NOR DNA. NOR measures are rRNA gene copy numbers in nucleolar organizing regions, where 4x,63/16 indicates copy excess (4x) for DNA over Asm, and total DNA/Asm amounts (Mb).

| SppID | ..... DNA Measures ..... |  |  |  |  |  | NOR_D/A | DNA_SRA_ID | Biosample |
| --- | --- | --- | --- | --- | --- | --- | --- | --- | --- |
|  | Cym | Gnodes | GSKap | GSKgs | GSKfi | Asmlen |  |  |  |
| A Arath | 157 | 146 | 150 | 137 | 139 | 134 | 10x,8-9/0.81 | ERR7919757 | Ara.th. Col-0 |
| T Tomato | 960 | 950 | 940 | 880 | 880 | 830 | 4x,63/16 | ERR13372748,9 | Sol.ly. heinz |
| M Maize | 2660 | 2630 | 2640 | 2440 | 2390 | 2200 | 8x,44/5.3 | ERR13381137 | Zea mays B73 |
| F Finch | 1220 | 1240 | 1120 | 1070 | 1440 | 1140 | 6x,1.6/0.26 | SRR32887368,9 | Tae.gut. bTaeGut7 |
| Z zFish | 1550 | 1550 | 1630 | 1470 | 1640 | 1450 | 7x,0.61/0.09 | ERR12118113,4 | Dan.rer. Tubingen |
| SppID | Cym | Gnodes | GSKap | GSKgs | GSKfi | Asmlen | NOR_D/A% of DNA - Asm |  |  |
| A Arath | .. | -7% | -4% | -13% | -11% | -15% | 71% |  |  |
| T Tomato | .. | -1% | -2% | -8% | -8% | -14% | 40% |  |  |
| M Maize | .. | -1% | -1% | -8% | -10% | -17% | 9% |  |  |
| F Finch | .. | +2% | -8% | -13% | +18% | -6% | 1% |  |  |
| Z zFish | .. | 0% | +5% | -5% | +6% | -6% | 0.5% |  |  |

Notes: GSK have non-nuclear contaminants removed (A contam=11mb, M contam=2mb) as measured by Gnodes, or from decontam readset (T, contam=62mb chloroplast+mito); Finch is zebra finch of 2025 from PRJNA1241433, ONT.10R and other DNA. ZFish is zebrafish, ONT.R10 DNA of PRJEB53713, corrected for 1% bacterial contam.; DNA for A,T,M of Bioproject PRJEB49840; DNA measures as in Table DC5. See Table DC1 for Cym, DNA data citations.

The rRNA content is measured and reported in nucleolar organizing regions (NOR) column of Table DC6. This is a simple and reliable measure of duplicated genome content that can be applied to all plant and animal genomes. In these samples the NOR spans are under-assembled, by 6 to 10 fold, accounting for 71% of Cym - Asm discrepancy of *Ara. thaliana* (see also Igolkina et al 2025, on Arath genome contents discrepancies), down to 1% in bird and fish. Though these later are fractions of the animal genomes, the unassembled rRNA genes are indicators of small, unassembled genome portions of biological significance. Other duplicated gene arrays are likely subject to similar mis-assembly problems. Assembly of these ONT R10 data, reported in detail Table DC7, recovers genome sizes close to Cym and DNA measures, when all assembly primary contigs are retained.

Figure DC2 is a graphic representation of the quality of DNA measures agreement with cytometry, for each population of three species population sets: *Zea may*s inbred lines, *Ara. thaliana* regional populations, and *Ara. arenosa* altitude populations. These plot DNA measure difference from Cym for each population, for DNA measures of Gnodes, GSK-GenomeScope, and population assembly sizes (excluding *Ara. arenosa*). Here the Gnodes differences are not significant (i.e. near zero), while GSKgs and Assembly are both below Cym and trend downward for increasing Cym measured genome sizes.

**Figure DC2.**
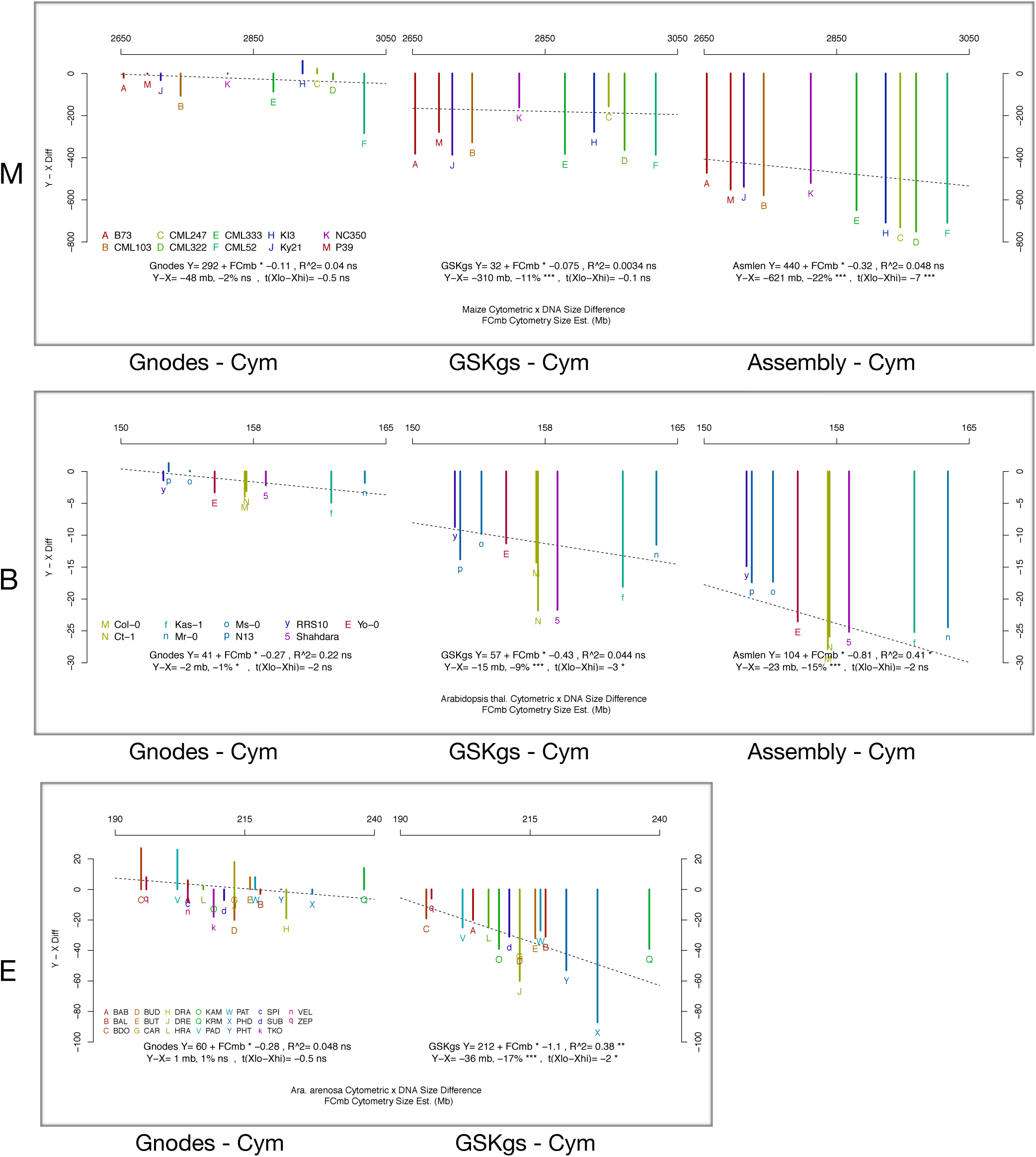
Differences of DNA size methods from Cytometry sizes, plotted for species populations: **M** Maize isolines, **B** *Ara. thaliana* regions, **E** *Ara. arenosa* altitude populations. DNA measure methods are Gnodes, GSK GenomeScope, and Assembly, left to right. X-axis is Cym size in megabases, Y-axis is difference from Cym size, negative but for Gnodes. Regression line and difference t-tests are listed, and population labels A to q are in lower left legend. These plots show details of quality summary of Table DC5.

**Figure DC3.**
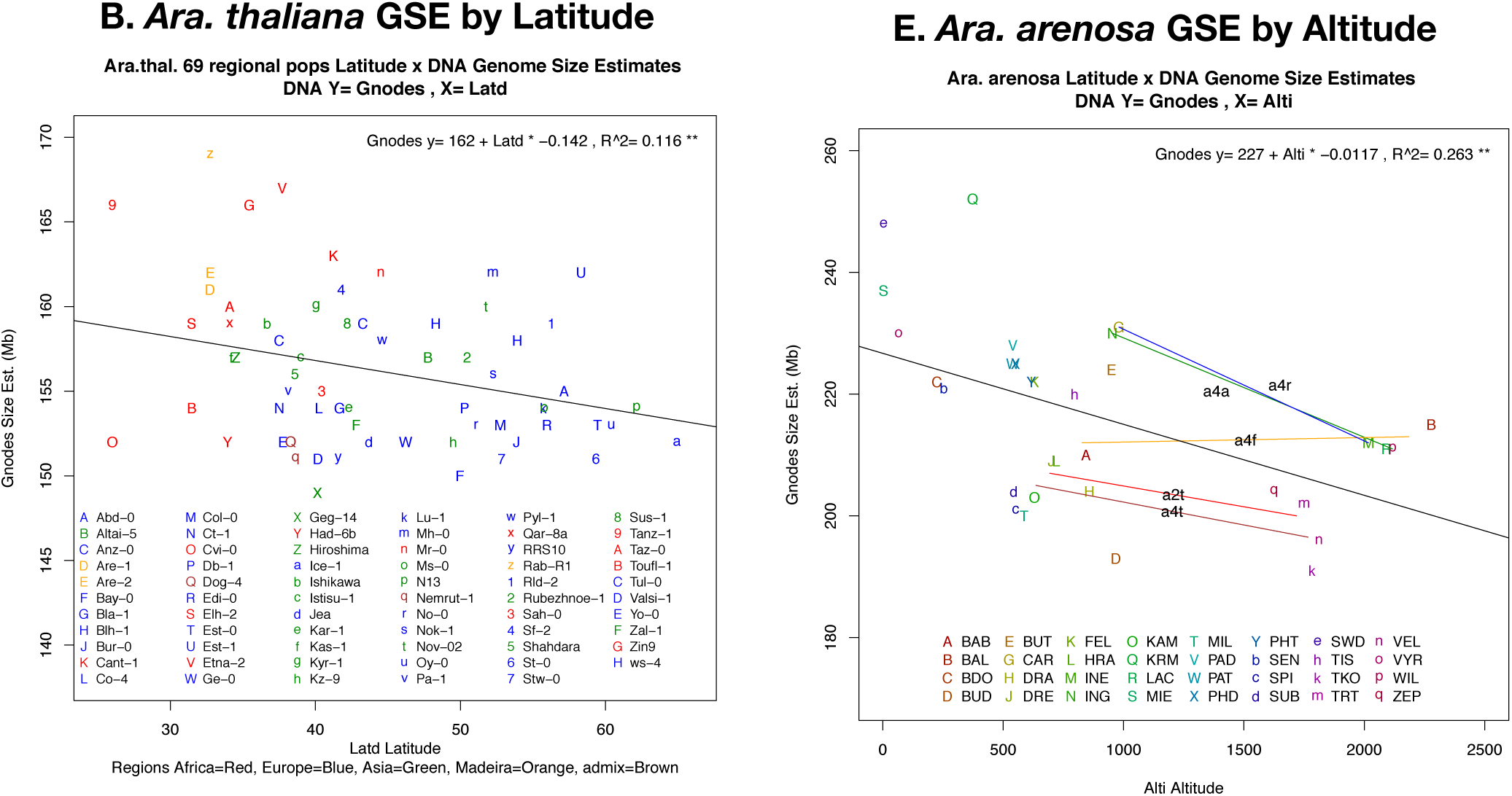
Altitude and Latitude effects for population genome sizes measured from DNA, for **(B)** *Ara. thaliana* regional populations by latitude, and **(E)** *Ara. arenosa* populations by altitude. For E, 5 locales with nearby alpine (high) and foothill (low) populations are shown by short lines that match the general cline of genome reduction at higher altitudes, in 4 of 5 cases. Y-axis is genome size measured with Gnodes, X-axis is altitude (meters) or latitude (degrees North). Regressions of GSE on these are significant; population labels A to q are in lower legend. These plots show details of Table DC4.

## Results in detail

When this project first examined DNA types in 2022, I located two projects of *Arabidopsis* ecotype populations that published Cym measures, and long and short DNA samples. These offered a gradation of putative population genome changes, measurable in two distinct ways, with comparable data of one species. Population variation offers a more controlled sample than genome measures across species. As summarized in Gnodes#2 Table T2.A, the difference measured for long - short reads is insignificant.

Further published Cym and DNA data was available for Maize inbred lines, where Cym measured a larger range in genome sizes (2,000 Mb to 3,000 Mb). One study (Hufford et al., 2021) of corn isolines provides short and long reads, and their assemblies, for a balanced, same-project data set. Gnodes measured a significant long-short read discrepancy: 334 Mb or 13% of genome sizes (Gnodes#2 Table T2.B). This also showed a close correlation of Cym and long-DNA measures, with isoline differences in the large maize transposon content. In these data sets there were, however, some possible differences between Cym and lr/sr DNA measures, but this data looked insufficient to resolve those differences.

### Biases in DNA sequencing methods lead to measurement errors

One of the important variables in measuring DNA for genome sizes is bias, or un-even/non-random coverage depth over the genome from sequencing technology and methods. The equation Genome_size = Shotgun_Read_Bases / Coverage_Depth assumes random, even coverage depth over a genome. This assumption is at the basis of genome-mapped and k-mer calculations, yet it is known to be violated by various methods of DNA sequencing. Early PCR-amplified DNA used for Illumina sequencing had large biases related to repetitive DNA GC-content; later “PCR-free” labeled methods, still using PCR or related amplification can have biases of non-random amplification of genome contents. Oxford Nanopore Technology (ONT) avoids PCR amplification, and thus may be better suited to genome measurement (Lerminiaux et al. 2023; Oxford Nanopore Technology document 2024).

Bias measures of genome content coverage are examined for the At model plant, three standard plant genomes, and recent zebra finch genome assembly and DNA, as shown in Figures DC4A (Arath), B (plants), Z (finch). In these examples, Illumina real and simulated biases reduce the apparent repeat contents of genomes, and are noticeable in the coverage graphs. This result, of ONT method having the least bias, was noted in analyses given in Gnodes#2 paper. The haploid sex chromosomes of female zebra finch contribute another bias in genome measurement, one than can be corrected. Effects of biases on total genome size measures depend of course on amounts of non-random coverage, but those are difficult to measure. With the annotated-assembly-mapped method of Gnodes, biases in major content types can be detected and partly corrected, under some assumptions (e.g. that the chromosome assembly and DNA mappings are mostly accurate). For the zebra finch example, these assumptions appear to hold; a result is that an unbiased ONT.R10 read set for the female can be corrected for haploid sex chromosomes (10% of genome), with a resulting measure that matches exceeds assembly size by ∼6%, and matches the cytometric size.

**Figure DC4A.**
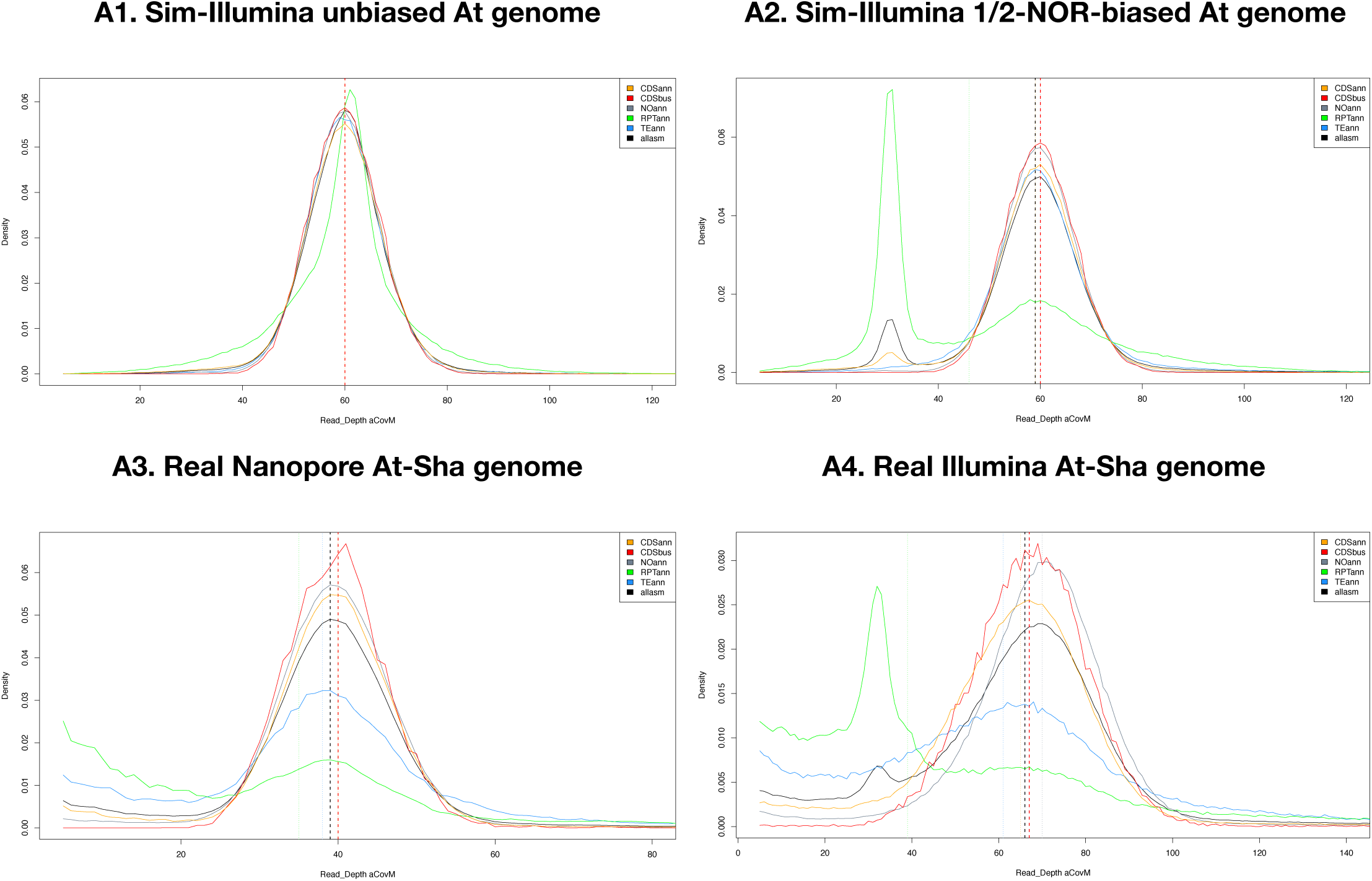
Biases in DNA shotgun reads shown in coverage depth graphs of *Ara. thaliana* genome major contents. Simulated DNA from complete *At* genome (A1), and (A2) biased by removal of 50% of NOR repetitive spans, 3% of genome. Real DNA reads of *At* ecotype Sha, for ONT long reads (A3, SRAid ERR11436646) and Illumina short (A4, SRAid ERR11436067), that shows a low-cover bias in repeats. Major contents are CDSann (orange), CDSbus (unique-conserved, in red), NOann (gray), RPTann (green), TEann (blue), and allasm (black). Vertical bars are at medians. An unbiased read set has similar coverage density for all contents; bias peaks of lower coverage are toward left.

The simulated bias (A2) was produced by randomly removing 50% of sim-reads in the NOR spans of Chr2,4, or 4.5 mb of the 9 mb NOR sequence, as assembled by [ref, GenBank IDs OR453402.1, OR453401.1]. The resulting coverage graph A2, by itself, can be interpreted two ways: 1. bias against repetitive regions in read amplifcation, or 2. a population genome that has 50% reduced NOR spans. A read set simulated from a genome assembly with 1/2 NOR spans, but without sim-bias read removal, has a coverage graph like A2. However the real At-Shahdara population genome of A3 excludes that second case: as measured with Nanopore reads, that a low-coverage peak doesn’t exist, and the genome size estimate is higher, matching Cym, versus the Illumina (A4) read set for this population.

**Figure DC4B.**
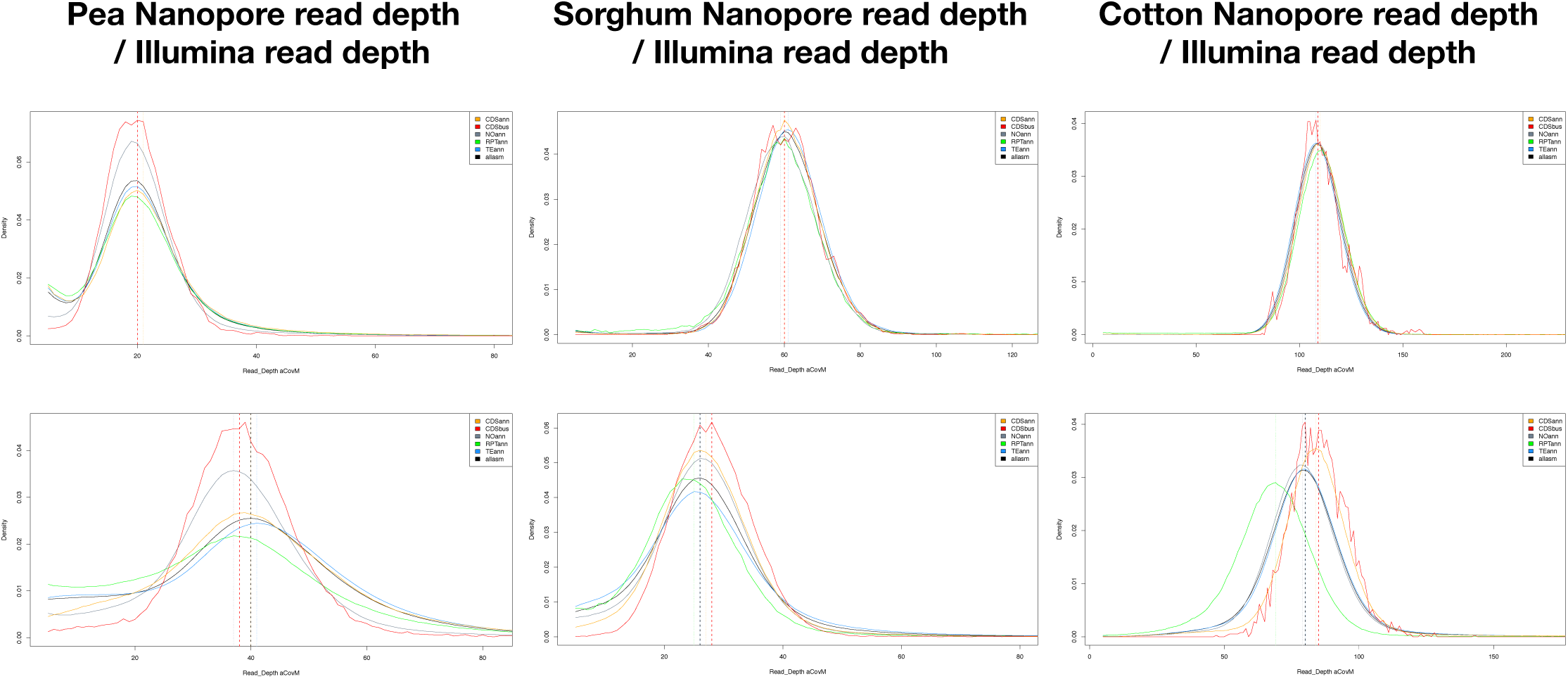
Biases in DNA read sets shown in coverage graphs for genome major contents, for three plant species pea, sorghum and cotton, with Illumina and Oxford Nanopore DNA. Major contents are as Figure DC4A. An unbiased read set has similar coverage density for all contents; bias peaks of lower coverage are toward left. SRAids are Pea ONT ERR9980782t97 & Illu SRR31015884, Sorghum ONT SRR26882277 & Illu SRR24011165, Cotton ONT SRR24575697 & Illu SRR24575699.

An extreme case of GC bias in DNA reads is the valuable but un-replicated work (Table DC2 1/A, Long et al 2013) on At ecotype lines that measured both Cym and DNA contents for 100s of lines in a geographic range, finding a cline in genome sizes consistent between Cym and DNA. The GC bias of this Illumina DNA is 6%, which obscures measures of ecotype genome variation, as this amplification bias differentially affects CDS, transposons, rRNA and other duplications. Re-examination of this old DNA data with Gnodes does recover the differences in genome sizes between N-S Sweden populations (Figure DC5), and DNA x Cym relation is significantly not different from 1:1, though these are weaker relations than for more recent DNA data.

Figure DC4z shows DNA coverage depth plots of zebra finch, female and male, from Gnodes measures of recent 2025 assembly and DNA sets of the Vert. Genome Project. ONT.R10 Simplex technology is designed to produce unbiased, un-amplified DNA. PacBio HiFi long reads and Illumina short reads are drawn from amplified DNA, in ways that can bias against duplications. The HiFi and Illumina depth plots of DC4z show repeat content biases, similar to low-skewed coverage for plant repeats from Illumina DNA (Figure DC4). HiFi is produced with a computation step of low-fi read averaging, which may also bias against highly similar duplications. This example confirms and extends results of Gnodes#2, that found ONT technology to be least biased. One ONT R10 Duplex data set, for Zea mays, showed bias in a large reduction of rRNA copy numbers (Gnodes #2), that may be a result of errors in early software to convert Simplex (2 strands of DNA) to Duplex (average of double stranded DNA), as the read use statistics for ONT Duplex for maize showed a large portion were discarded, versus Simplex. As measured here, the ONT.R10 Simplex of maize fully recovers the duplicated genome contents (Table DC6).

**Figure DC4Z.**
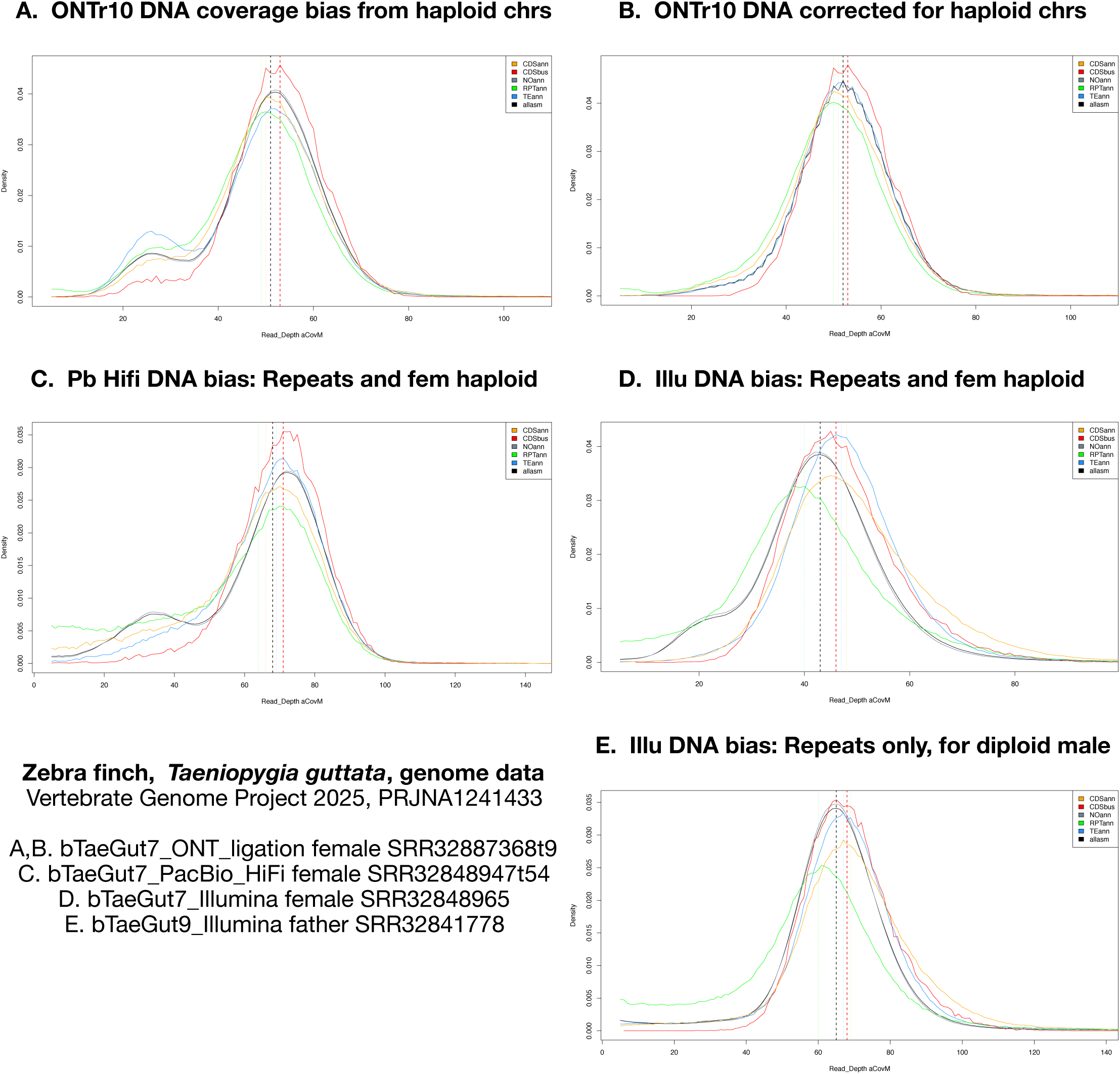
Bias in coverage depth from haploid sex chromosomes in ONT.R10 DNA of female zebra finch (A,B), and biases in repeats and duplicated contents for PacBio HiFi (C) and Illumina DNA coverage (D female, E male). The correction of (A) for 1/2 ploidy is shown in (B). Female bird haploid chromosomes W, Z are 10% of genome, shown as smaller peaks at ∼25 versus main ∼50 peak depth (A). Male has diploid Z, no W, thus no 1/2 depth peak, but shows Illu bias against RPT content. Genome total coverage and major content types are as Figure DC4, see also Table DC6.

### Population genomics details of DNA measures and Cytometry

Details of DNA by Cytometric measures for Arabidopsis populations are given here. Figures DC5 show the DNA x Cym relations for early At1K data (Long et al 2013), which have weak but statistically significant correlations. The weakness is primarily due to bias in this early generation short-read DNA. Figures DC6 show these DNA by Cym relations for more recent DNA of *Ara. thaliana* of 69 regional ecotype populations (Lian et al 2024). The Cym values for these populations are drawn from same earlier At1K work by matching population IDs, using an untested assumption that these populations retain the same genome sizes.

**Figure DC5.**
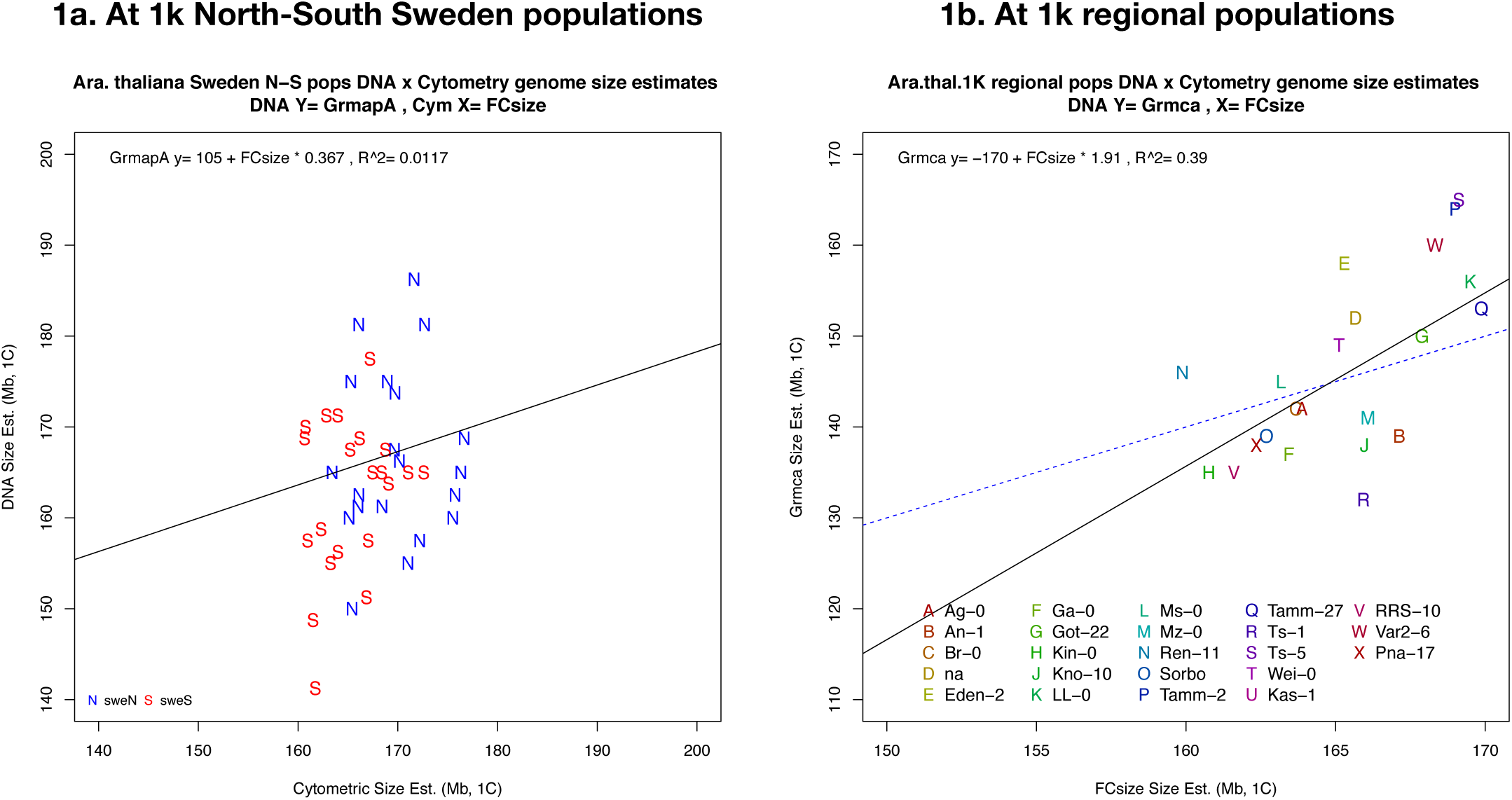
DNA x Cym relation in *Ara.thaliana* of Sweden with North-South populations size effect (1a), and *Ara.th.* regional ecotype populations (1b).

**Figure DC6.**
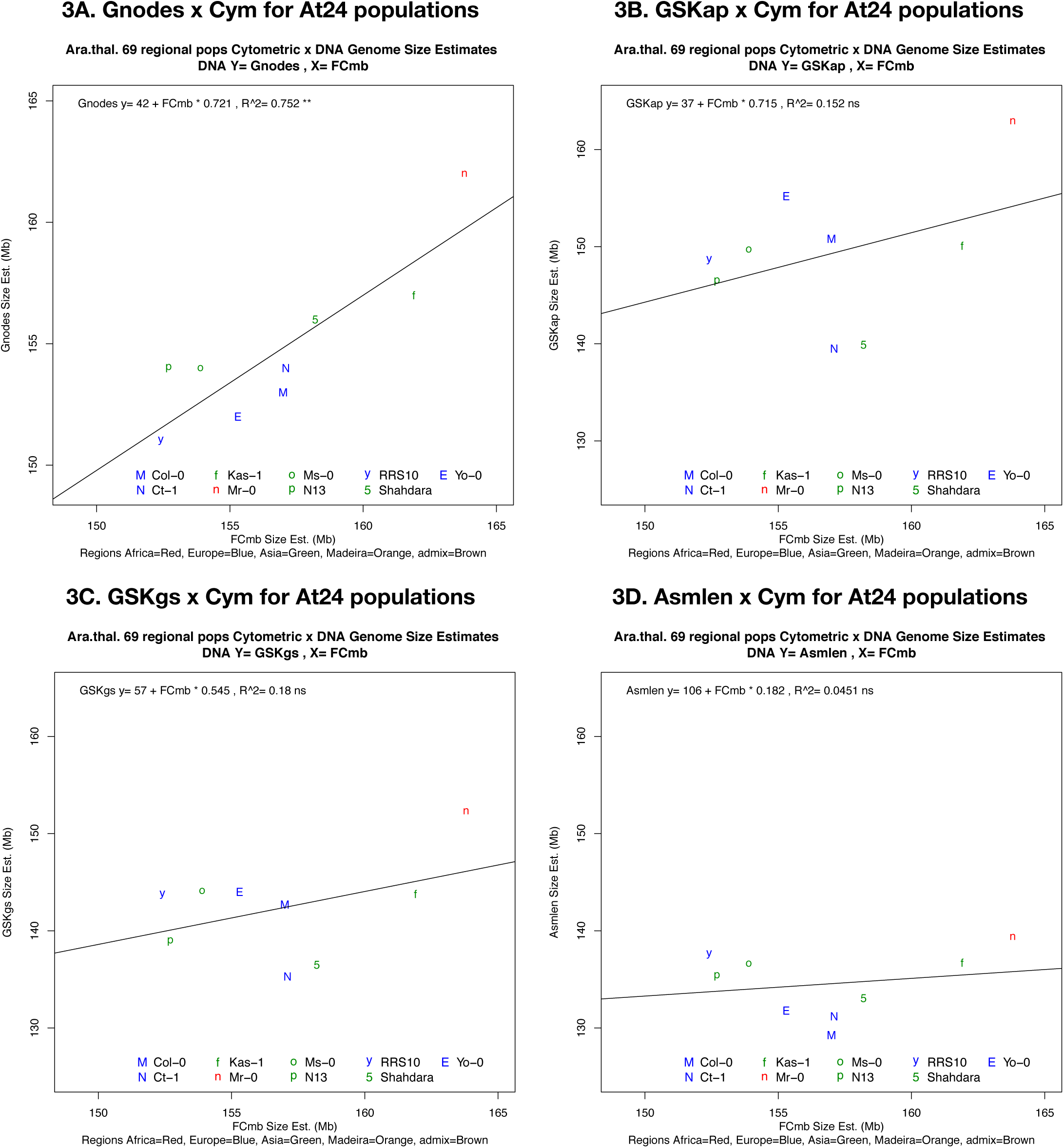
DNA x Cym relation in *Ara. thaliana* of 69 regional ecotype populations (2024 ref). DNA measures of Gnodes (3A), Kmer-peak (GSKap, 3B), Kmer-GenomeScope (GSKgs, 3C), and Assembly (Asmlen, 3D) are shown. DNA x Cym regression is significant for Gnodes, but not the other 3 DNA methods. This is a subset with Cym values from the earlier At-1k (2011) study. The latitudinal effect in these populations is shown in Figure DC3.B. Plot symbol colors indicate the major regions of this study.

**Figure DC7.**
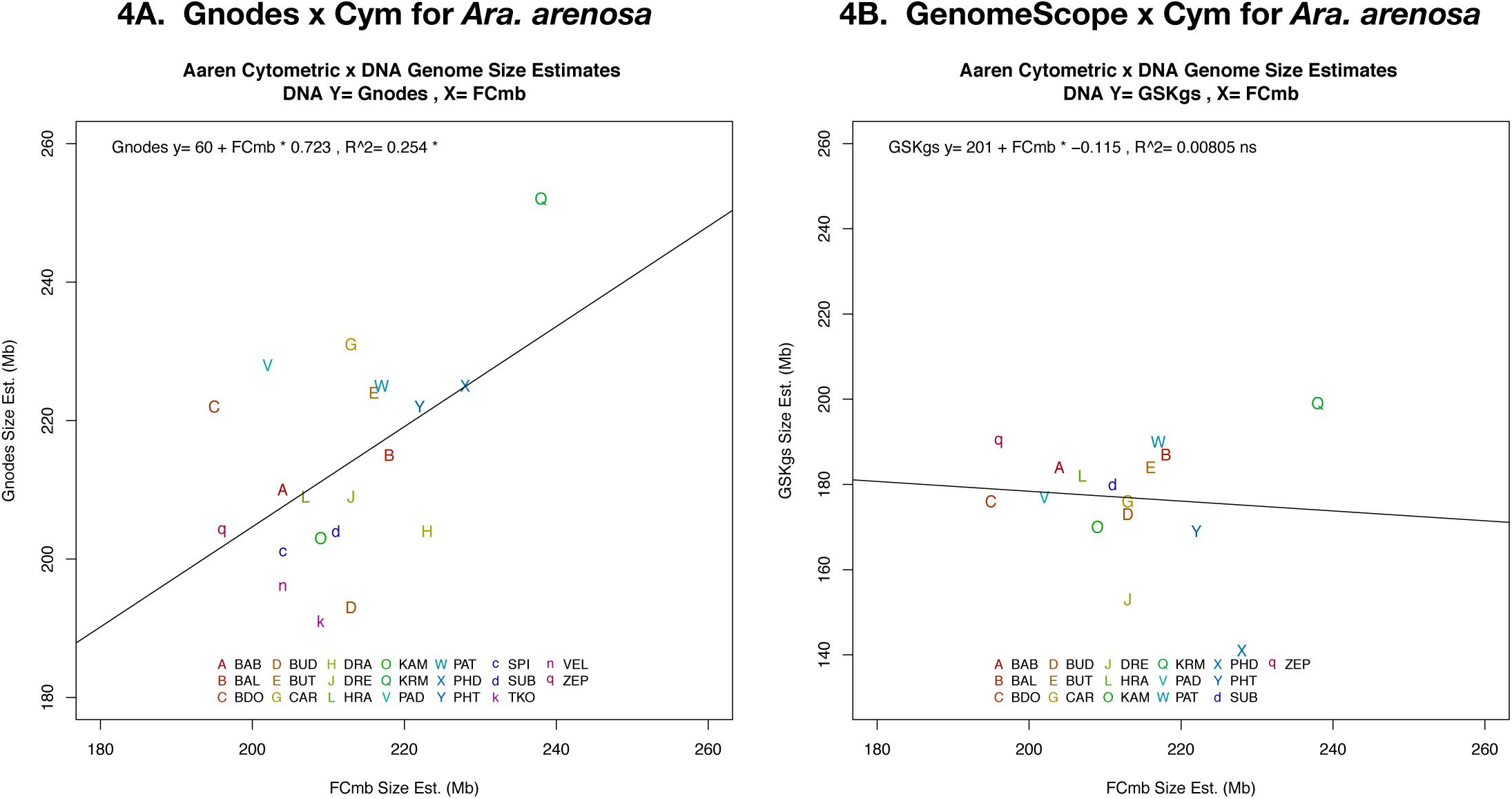
DNA x Cym relation in *Ara. arenosa* of altitudinal populations. DNA measures of Gnodes (4A), and Kmer-GenomeScope (GSKgs, 4B) are shown. DNA x Cym regression is significant for Gnodes, but not for GenomeScope. The altitude effect in these populations is shown in Figure DC3.E.

**Figure of Table DC7.**
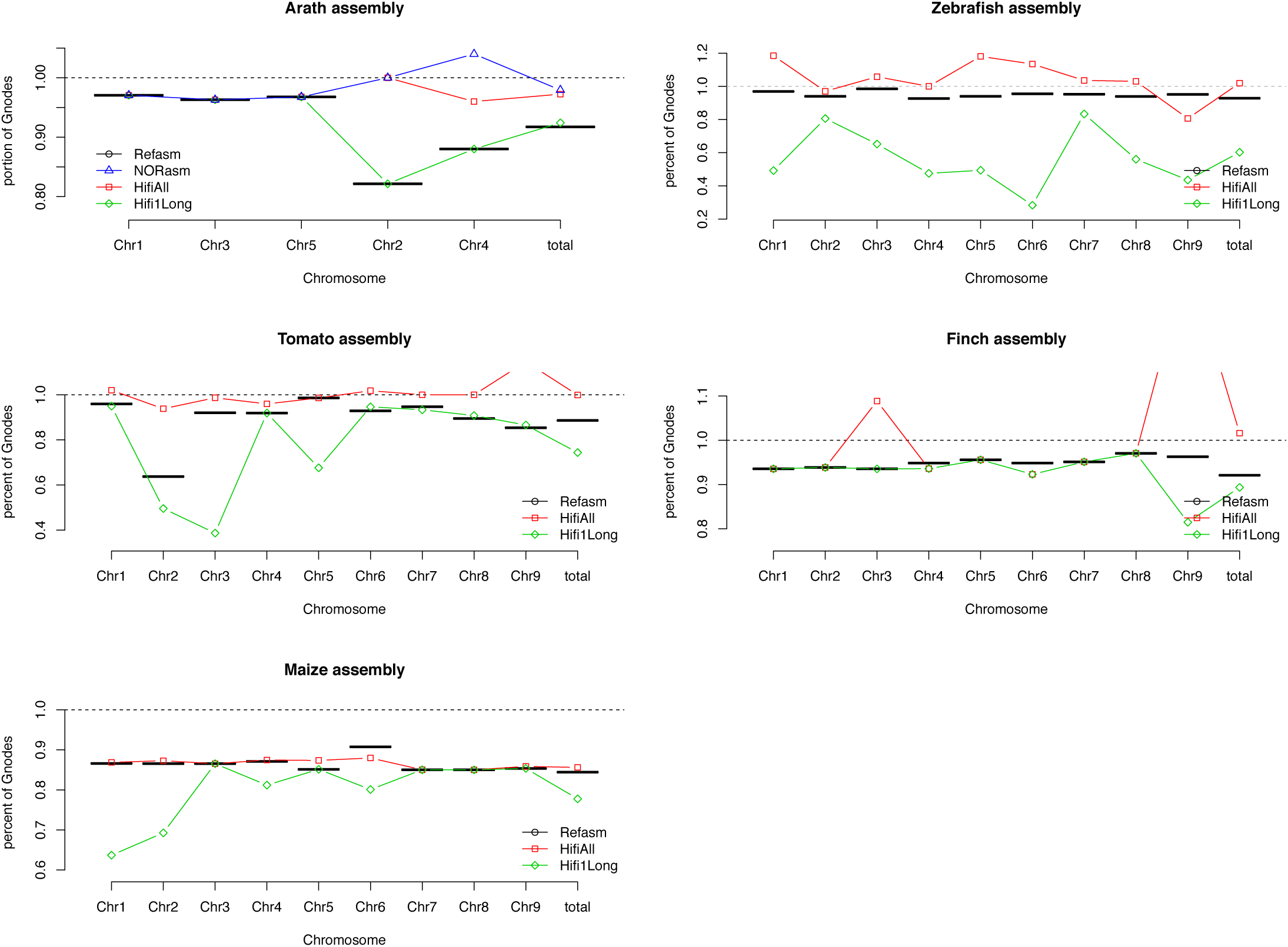
Chromosome sizes for ONT R10 Simplex of Arath (A), Tomato (T), Zebrafish (Z), Finch (F) and Maize (M), as measured from DNA with Gnodes, assembled with hifiasm of same DNA, and of reference assemblies. Plots show sizes relative to Gnodes (dotted line at 1), for the subset of Chr1 to 9. The longest hifiasm contig (Hifi1Long), and all hifiasm primary contigs (HifiAll) aligned to reference are shown. NORasm of Arath is a reference with NOR assemblies appended to Chr2,4 where they belong. Details are in gnodes4tables/gnd25cymdna_ontr10asm_tab.txt, and Table DC6.

**Table DC7.**
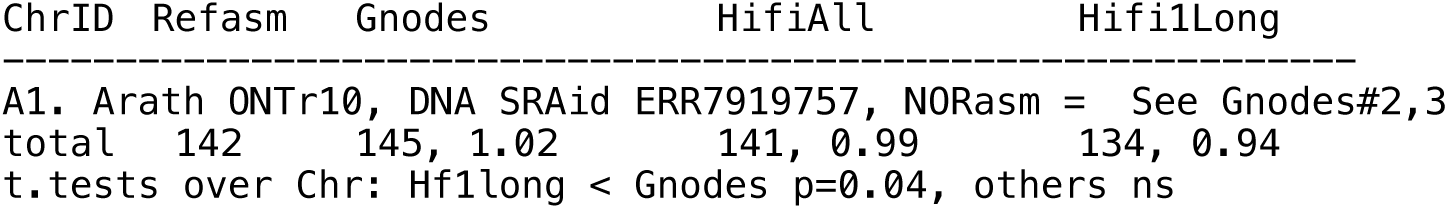
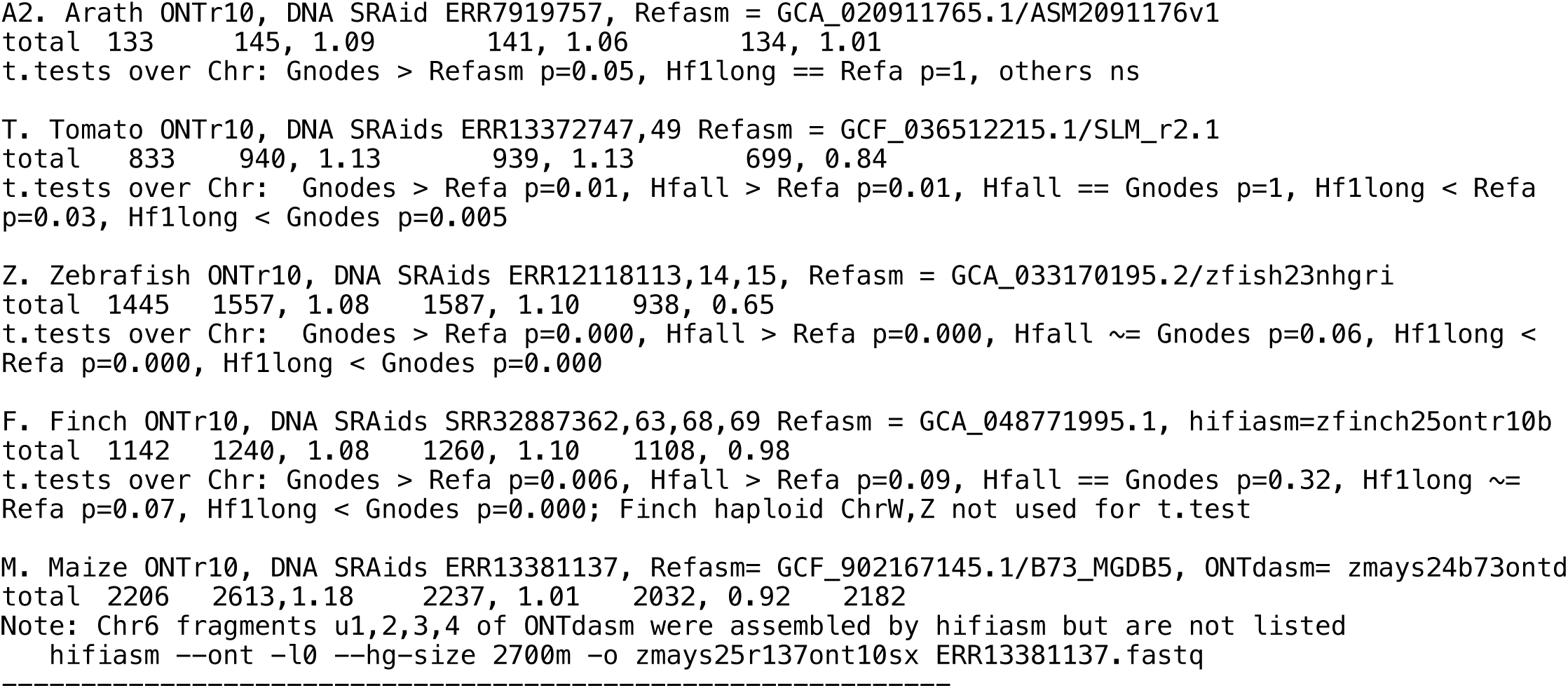
totals only. Chromosome sizes for ONT R10 Simplex of Arath (A1,A2), Tomato (T), Zebrafish (Z), Finch (F) and Maize (M), as measured from DNA with Gnodes, from hifiasm of same DNA, and from reference assemblies. Sizes in megabases, and ratio of size to ref assembly are given. Single longest hifiasm contig, and all hifiasm contigs are given. Reference assemblies for Arath are A1. NOR-extended “complete” assembly, 142 Mb, and A2. published reference T2T assembly, 133 Mb [GenBank GCA_020911765.1 of 2022]; Tomato published T2T assembly [SLM_r2.1, RefSeq GCF_036512215.1 of 2024]. Gnodes and HifiasmAll measure about the same sizes for chromosomes. Hifi1Long assembles the majority of A and T chromosomes as one contig, but needs more contigs for the difficult NOR spans and other duplicated sections. For A2, Gnodes measured on a smaller ref asm results in same larger NOR Chr2,4 sizes; Hifiasm is not measured against ref asm. See Supplemental Table DC7 in gnodes4tables/ gnd25cymdna_ontr10asm_tab.txt, and Table DC6.

**Figure DC8.**
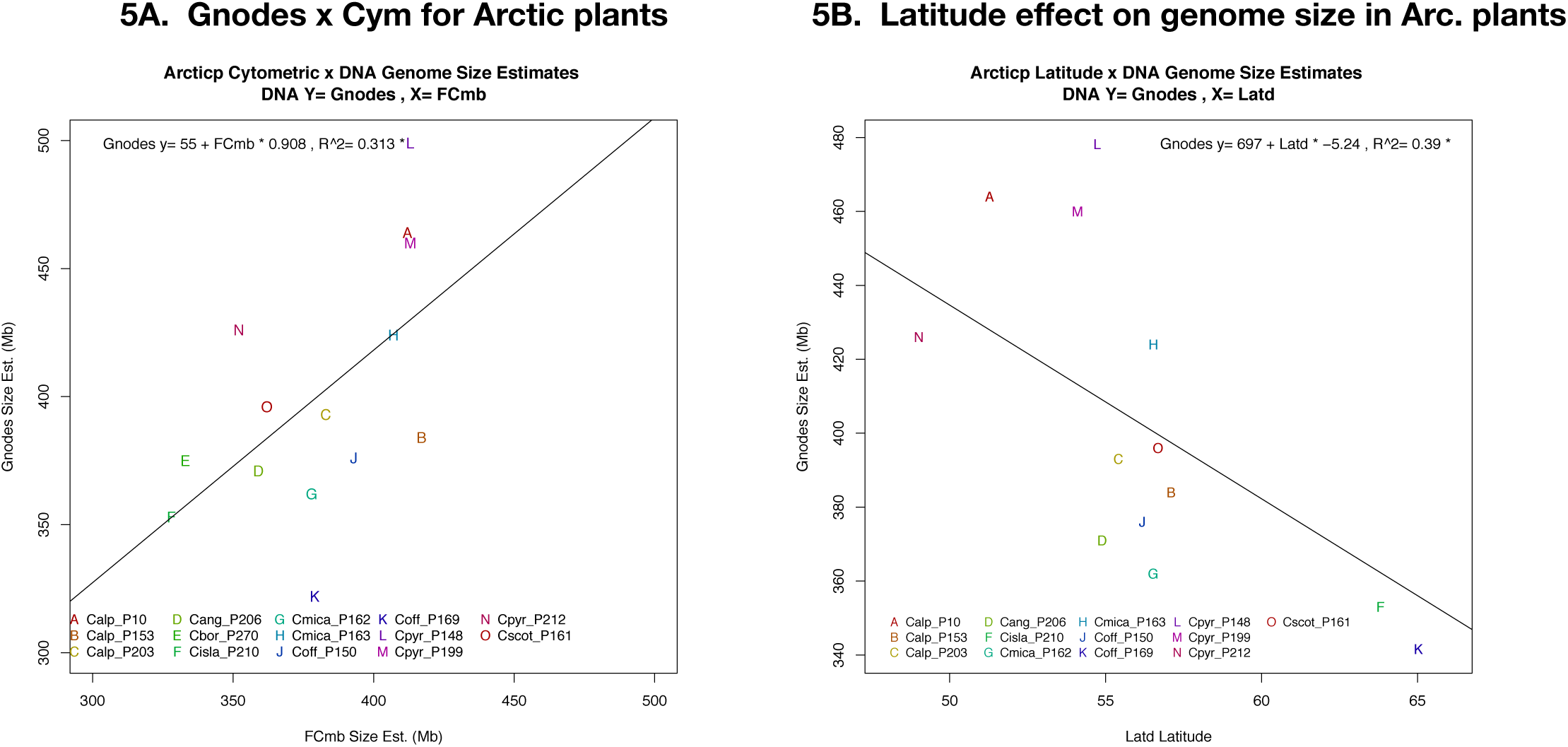
DNA x Cym relation in arctic plant populations of *Cochlearia* genus. DNA measures of Gnodes (5A), and latitudinal effect (5B) are shown, regressions are significant for both.

**Figure DC9.**
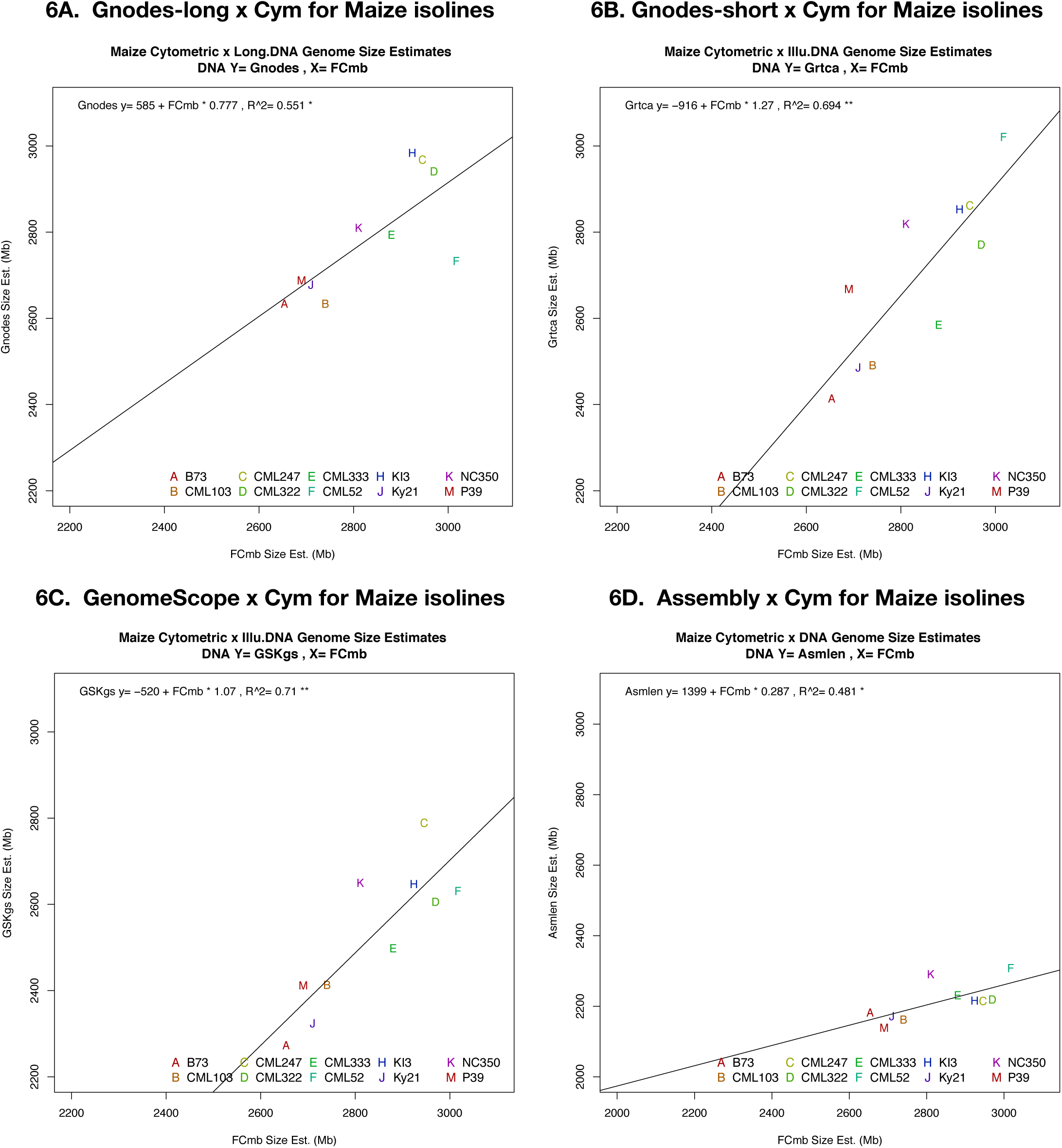
DNA x Cym relation in *Zea mays* inbred lines. DNA measures of Gnodes-long (6A), Gnodes-short (6B), Kmer-GenomeScope (GSKgs, 6C), and Assembly (Asmlen, 6D) are shown. All regressions are significant, but different: 6A of long-DNA is close to 1:1; 6B,C of short DNA are similar; 6D Assembly slope is ∼1/4 of Cym.

**Figure DC10.**
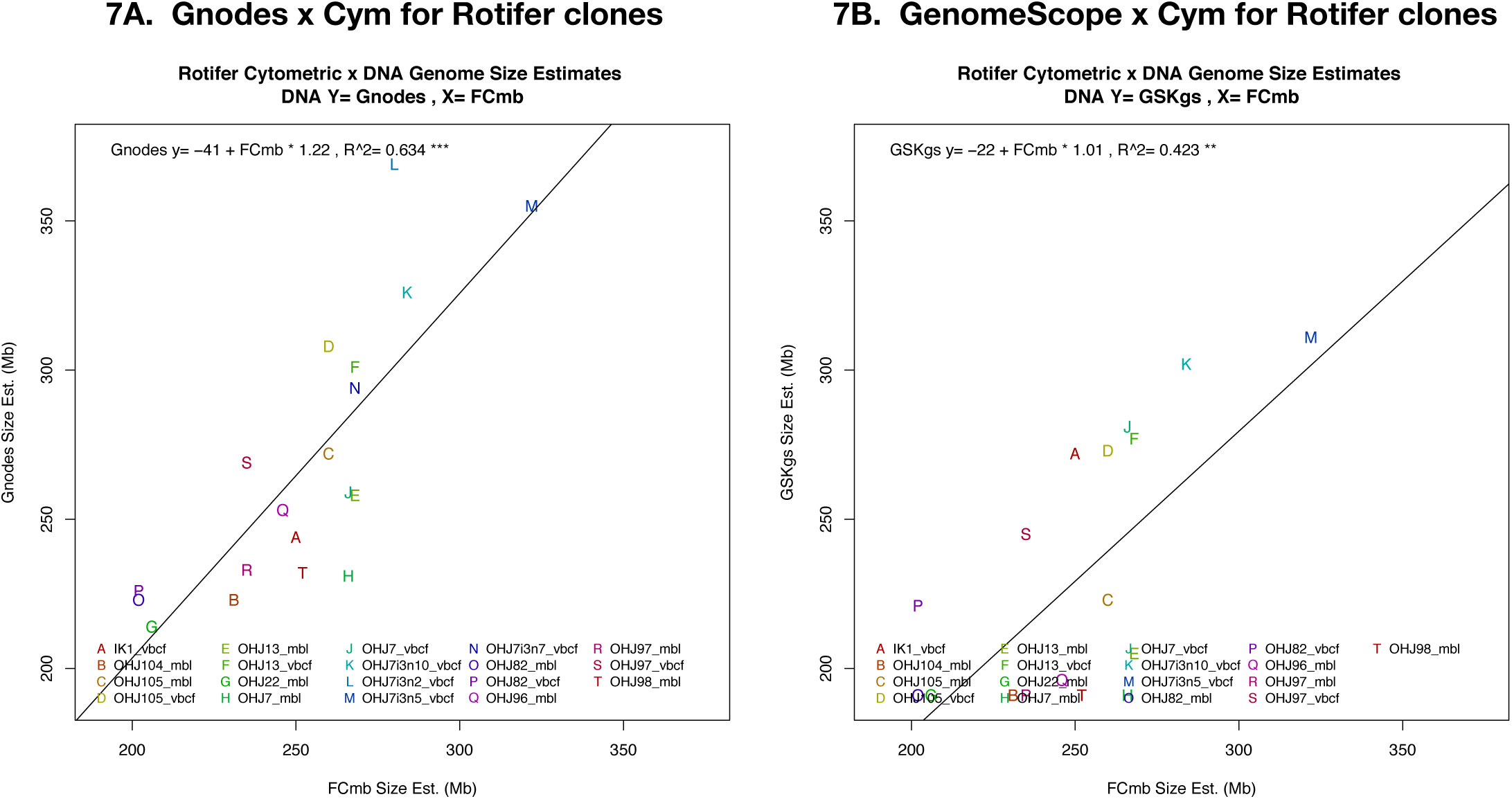
DNA x Cym relation in rotifer clones. DNA measures of Gnodes (7A), and Kmer-GenomeScope (GSKgs, 7B) are shown. Regressions are significant for both.

### DNA assembly can agree with Cytometry and DNA measures

The samples of ONT R10 Simplex DNA for five species agree with cytometry measures of genome sizes, in Table DC6. Do assemblies of that DNA fail to reach the same levels? For recently published assemblies, including some using the more accurate ONT R10 data, they remain 5%-15% below these sizes. However, the hifiasm software has recently been improved to assemble ONT R10 Simplex DNA [Cheng 2025], and results are in agreement with the Cym and DNA measures, when all assembly contigs are measured. Table DC7 lists the chromosome level assembly results of hifiasm-ont for Arath, Tomato, Maize, Finch and Zebrafish (data of Table DC6). The single longest hifiasm-assembled contig contains a majority of essentially complete chromosomes. However Arath Chrs 2,4 with large NOR spans are incomplete without added small primary contigs. For tomato, 5 of 12 are only partly assembled in one contig, the other assembled contigs fill in to the measured 940 Mb genome size, versus its recent reference assembly of 833 Mb. Maize ONTr10 DNA assembled with hifiasm did not reach the size measured in that DNA and by Cym. Finch hifi chromosomes are close to complete in single contigs, and similar in sizes to a recent reference assembly, but average 5-10% below Cym and DNA measures. Zebrafish assembly reaches the size measured in DNA and by Cym, but has a low portion of completed chromosomes, possibly because the DNA was not collected for assembly, but for other studies.

One likely reason for the discrepancy with published T2T assemblies is removal of extra assembled contigs that do not fit neatly into single chromosome spans, i.e. the tangles or bubbles from attempts to assembly long duplicated spans. This trade-off between simple, clean assemblies and complete, but fragmented ones is one that genomicists have wrestled with for 25 years. Though DNA data has improved much, this problem remains to be solved. Accurate measurement of genome sizes and contents is needed to indicate whether genome assemblies are missing contents; both cytometry and DNA-mapped measures appear to offer more accurate measures, when care is taken with methodology.

### Kmer DNA measures have basic flaws that underestimate genome sizes

Can lower size estimates from DNA by kmer methods be reconciled with those measured with cytometry and DNA-alignment by Gnodes (Table DC5, and results above)? Read depth plots of Figure DC11 are similar to those of GenomeScope and findGSE, which show peak read depths used to estimate total genome sizes. Both Error and Repeats portions of these kmer distributions contain large genomic DNA amounts. Repeats of DNA are added to GSK after depth is modelled on the main (homozygous) peak. However, the “Erroneous” distribution has a large amount of DNA unaccounted for by kmer models, and evidence indicates this is not erroneous for genome size measures. FindGSE and GenomeScope are similar in this aspect. CovEst and RESPECT kmer methods include rare kmers though it is unclear if this improves their GSE (see below).

**Figure DC11.**
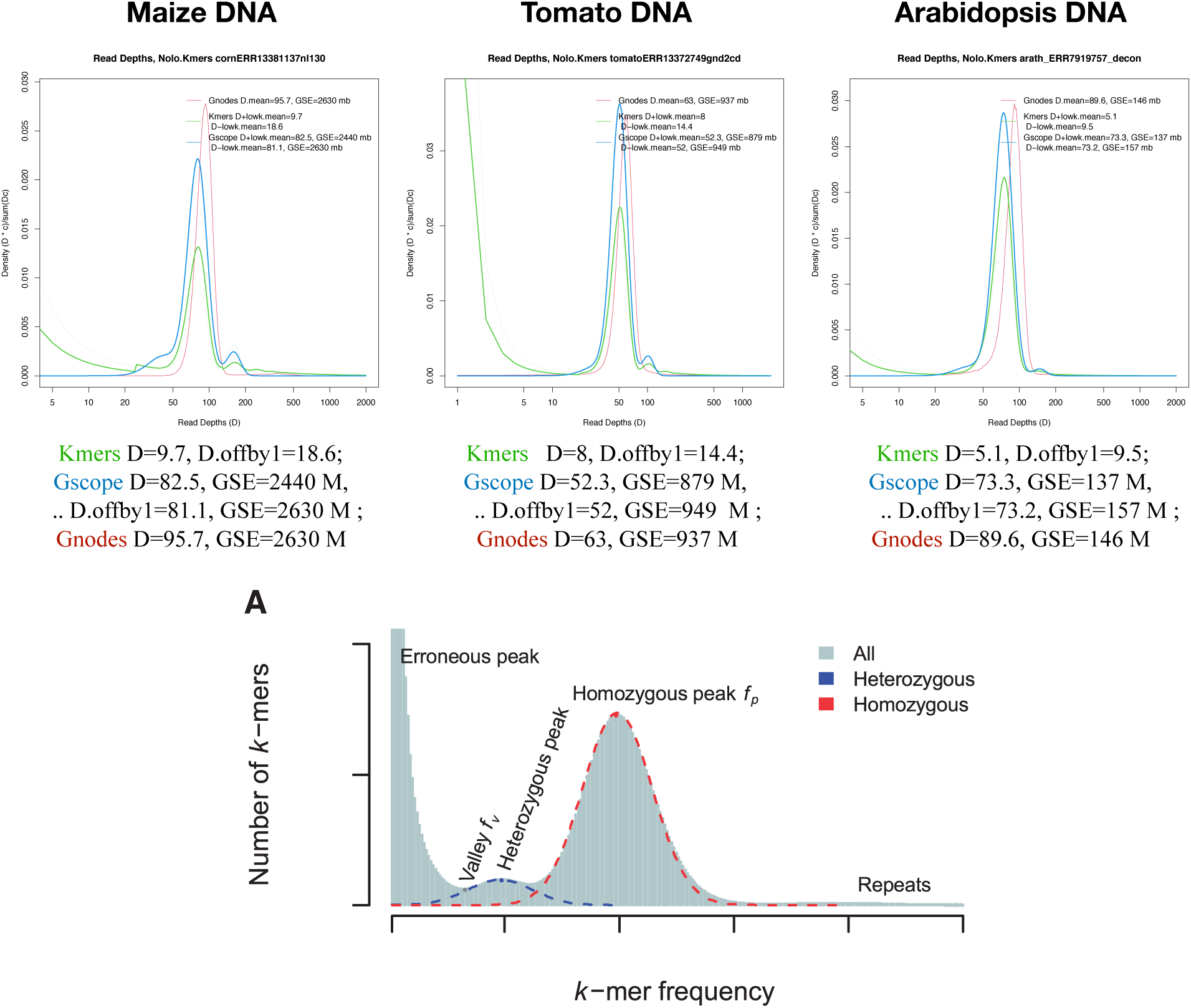
Read coverage depths, for Kmer histograms, GenomeScope model and Gnodes, from ONT R10 Simplex DNA mapped to nuclear genome assemblies of Maize, Tomato and Arabidopsis (see also Table DC6). These are density plots, x-axis is read depth (D) on log scale, y-axis is density of reads, (D * c count)/sum(D*c). The labeled Figure 1A of FindGSE paper [Sun et al 2017] shows three groups, Erroneous distribution (left), Homozygous distribution (center) and Repeats (right), that are all visible in DNA depth distributions of the 3 species. Values for Kmer histo., G.Scope model, and Gnodes are given as mean Depth (D) and GSE, of original kmer histogram (dotted lines), and with 50% of low-count “offby1” kmers shifted to the main distribution (solid lines). Note: D Depth of cover == C Cover depth

Mean depth given in Figure DC11 is the best measure of total *mass* of DNA read depth. Note that Kmer histogram mean D is very low, relative to GSE-K peak estimates, indicating a large mass of kmers in this “error” class. Supposing many of these rare kmers belong to the main mass for genome size measures, shifting 50% of these to the main distribution has noted effects.

The Gnodes distribution, equivalent in meaning, is measured not with kmer counts, but from a histogram based on read-alignment depth across genome assembly spans. The DNA from which the kmer fragments were taken all aligns to nuclear genome assemblies of the same DNA (chloroplast DNA is removed). This indicates that mers from these 20 kb average long reads, clearly part of genomic DNA, have some combination of base changes and indels that produce a large fraction of low-frequency mers. A portion of this ‘error’ group is accounted for as being off-by-one types of SNP/indel sequencing errors. For genome size measures, a nucleotide base value is not important, rather it is the location as a valid, unbiased position in a genome. Insertions and deletions, unless artifactually biased, are likewise not an error for size measurement. Discarding these low-copy kmers may introduce a bias in modeling a kmer distribution; a likely hypothesis is that these are drawn from the lower, less frequent end of the major density distribution.

If a portion of ‘off-by-1’ kmers are reclassified as belonging to the major density that is modelled by Kmer methods, then the density shifts (mean D in Fig. DC11), and kmer peak estimates are reduced a bit, thus recovering more genome DNA. For this ad hoc example, 50% of ‘error’ kmer mass is reallocated to the major density distribution, in proportion to density mass over that distribution (see methods for DC11 Example calculations). The results of corrected accounting of Error and Repeat kmers can be GSE-K estimates that are very close to Gnodes and Cytometry measured genome sizes (Fig. DC11, GScope D.offby1 values).

Further details of ‘off-by-1’ kmers were found by blast alignment to main kmers, collecting off-by-1 matches. These cause a downward shift in main kmer distribution, as shown by leftward shift in Figure DC11b. Unique 21-mers, with no other perfect matches in DNA, are a very large portion of these DNA samples. Most will match with 1 base change or indel to main-mers that form basis of C depth estimate by GSK methods, and will shift downward that C estimate, and upward the genome size estimate, using reasonable assumptions of their non-error distribution. GenomeScope calculations on corrected histograms are close to Cym size measures. Measurement details for this off-by-1 test are in methods section ‘DC11b Kmer-error off-by-1 analysis’.

**Figure DC11b.**
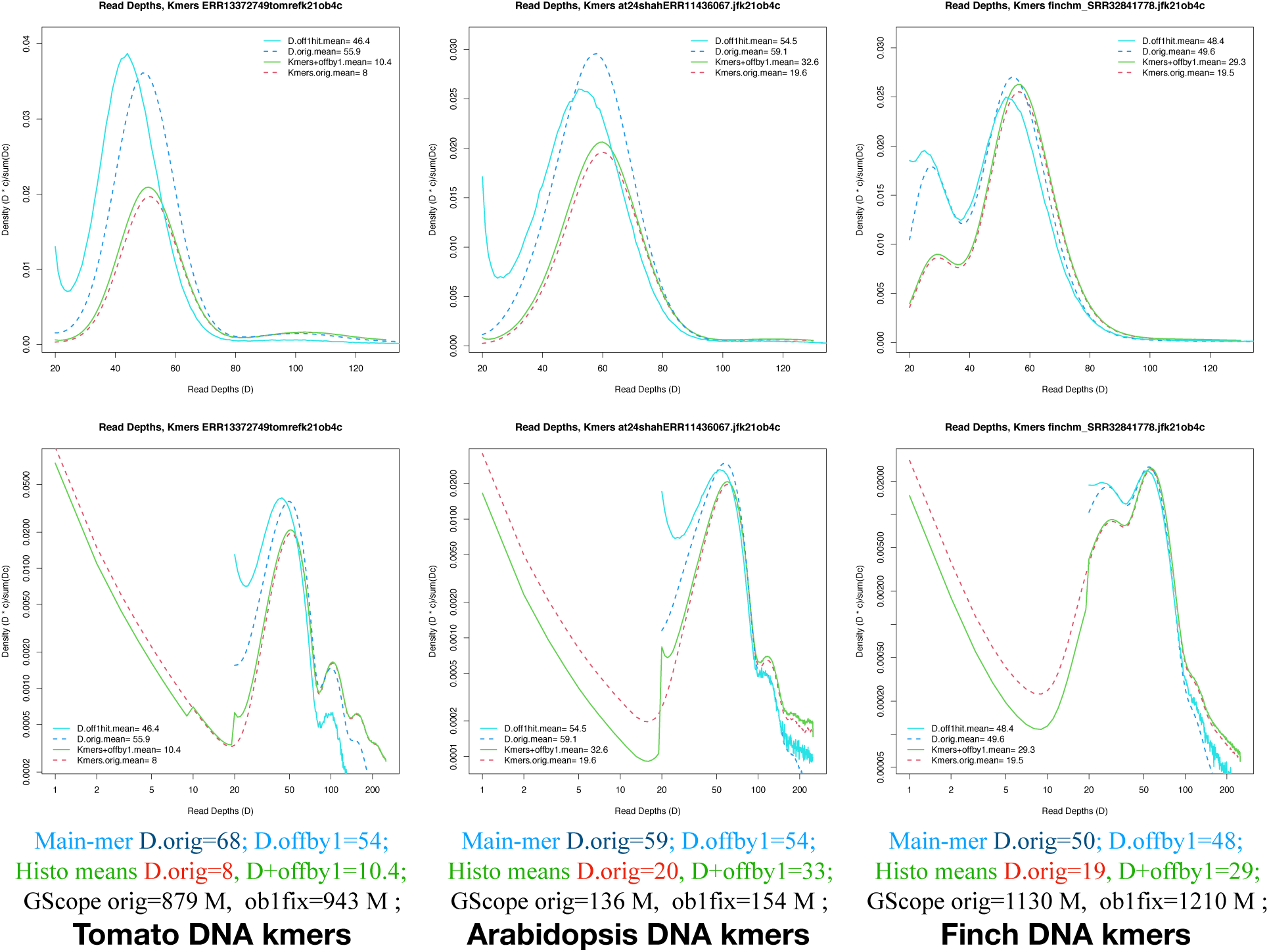
Test of kmer densities with off-by-1 error corrections, and uncorrected, for tomato ONT R10 DNA on left, Arabidopsis Illumina DNA middle, and finch Illumina DNA right. The plots show densities around main-kmer peak, for unique kmers matched to main kmers (solid lines), and the original main-mer density (dashed). Upper curves in blue are the densities of off-by-1 mers found in main-mer set, and unshifted main mers. Lower green and red curves show the full k-mer histogram density (y) per mer-frequency (x) that GSK methods use. Top plots are main-mer range (frequency 20-120), bottom plots are error+main range (1-250, log xy).

Can we tell if unmeasured rare k-mers have affected GSK estimates of population genomes? Yes, there are significant reductions in GSK with increasing unique k-mer density for some of the cases reported here, while for other cases, the low-count kmer effects are ambiguous (Fig. DC11c). Of the population sets with cytometry data (Tables DC1, DC2), only *Ara. arenosa* (E) had sufficient and suitable data for kmer analyses, and there is a strong negative correlation of unique kmer density and GSK size estimates, compared to Gnodes and cytometry measured sizes. *Ara. thaliana* of the 2024 population study (B) has a mixed result for Illumina DNA samples used for kmer analysis, strongly affected by laboratory effects (IlluOnt vs IlluPb labels of Fig DC11c p4). Recent population genome studies with rice (*Ozyra rufipogon* and *O. sativa,* rice pangenomes of Guo et al. 2025) provide PacBio HiFi DNA, which avoids some of the bias apparent with Illumina and can be analyzed with GSK methods, although no cytometry measures are available for these populations. These rice populations show significant negative correlation for unique kmer density and GSK size estimates (Fig DC11c p3).

**Figure DC11c.**
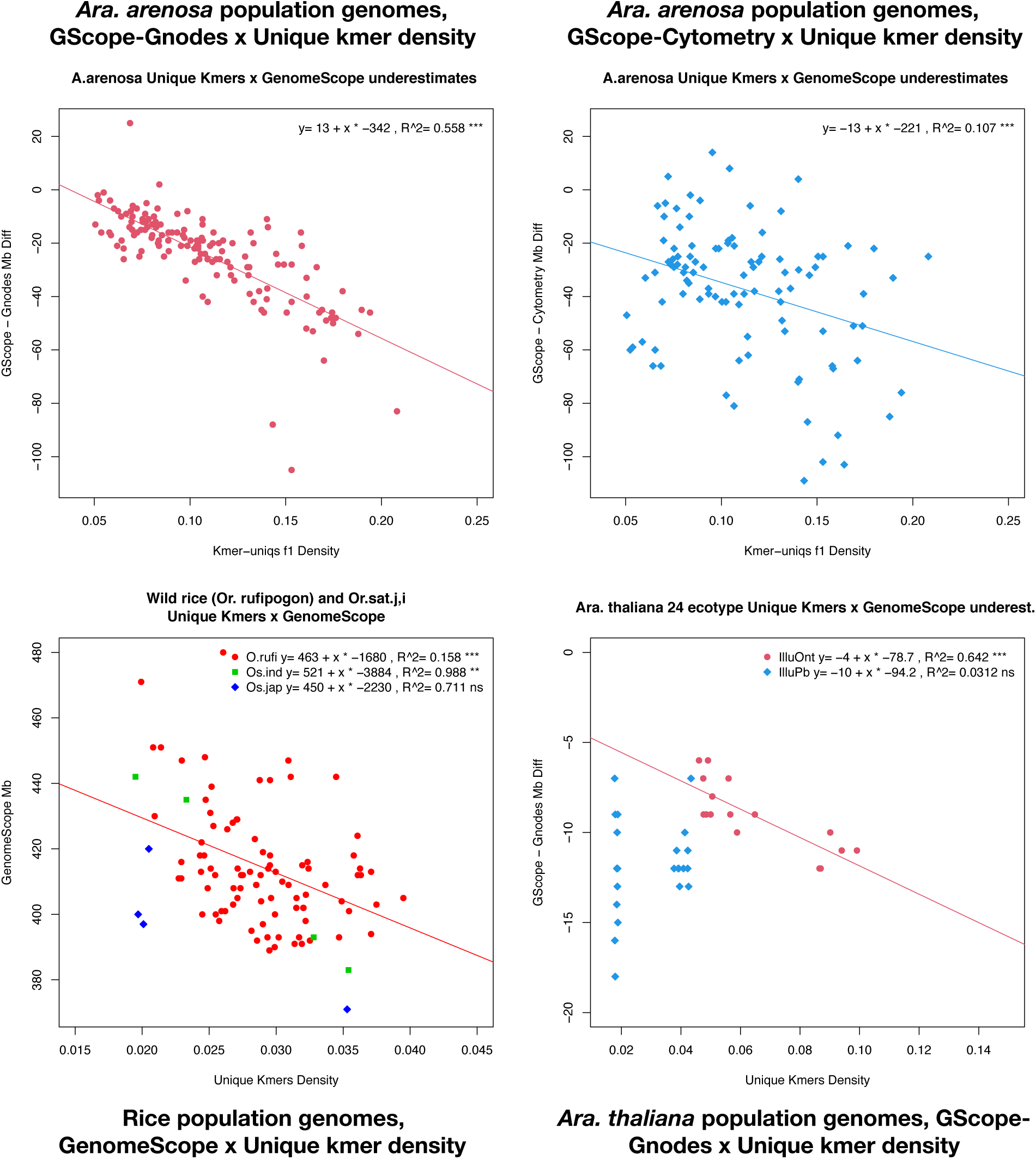
Unique kmer effects on GSK estimates by GenomeScope across population genomes, for *Ara. arenosa* (p1,2), *Ara. thaliana* (p4) and rice (p3, *Ozyra rufipogon* and *Oz. sativa* indica, japonica). X-axis is density of unique (f1) kmers, y-axes are GenomeScope GSE in megabases, minus Gnodes or Cytometry GSE for Ara. arenosa, A. thaliana. Regressions are given; the slope estimates, for 100*density (i.e. percentage f1) are megabases amounts that percentage increases in unique kmers will reduce the GSK estimates. E.g. a 1% increase in unique kmers for rice would reduce GSK by 20-30 Mb.

Maize isoline populations did not show a low-count kmer effect, however the Illumina data available for the 12 populations is biased which may obscure kmer frequency effects. The 21 rotifer clonal populations have Illumina DNA that also shows strong lab effects, where that of one lab has a non-significant trend to negative correlation for unique kmer density and GSK size estimates, the other lab DNA does not. GSK estimates for Illumina DNA of Arctic plant (N) populations all failed, due in part to low coverage depth. Only GenomeScope measures are analyzed here; findGSE or other GSK methods may have different results for rare kmer proportions. CovEst (Hozza, Vinar & Brejova 2015) explicitly includes unique and rare kmers in its “abundance spectra” modeling, but CovEST GSE’s are less accurate (both above and below Cym, GSK-A and other GSE-K methods, per several reports: Sarmashghi et al 2021, Gilbert 2022-23, Hesse 2023, ..) . RESPECT (Sarmashghi et al 2021) models GSE-K in a different way, including rare and overabundant k-mers. There are limited published results for this method, but Hesse (2023) reports it as the most accurate GSE-K method, with close match to Cym values, for 5 plant species. It seems likely that all GSK methods have difficulties accounting for the genomic contents of rare k-mers, as they all rely on perfect matches of imperfect mers to other genome data.

Repeat content is also a large fraction in the long right tail of read depth distribution, and accounts for a large portion of GSE. One author-correctable problem with kmer methods is that software will by default truncate the histogram used by GSE-K, removing high-copy repeats, thus producing erroneously low estimates. It is common for GSE-K software users to use these default settings. Gnodes #1, #3 (Gilbert 2022-23) documented under-estimates of genome size by GenomeScope and other kmer methods; this truncation cuts GSE-K estimates substantially (Arabidopsis: trunc 125 vs full 151 Mb, Daphnia: trunc 137 vs full 242 Mb; Gnodes #3, Table 3C).

Even when changed, very high copy kmers from repeats can be missed: in an *Arabidopsis* genomes publication (Lian et al. 2024), a high cut-off of 2 million still truncates the full distribution by 3%-5% of genome sequence. Current GenomeScope how-to-use documents do not advise against this default truncation of repeats in the kmer histogram. GenomeScope version 2 paper (Ranallo-Benavidez, Karon and Schatz 2020) does mention this in discussion, reversing v1 recommendations (Vurture et al 2017). Current findGSE README suggests ‘jellyfish histo -h 3000000’, but this still truncates very high copy genome repeats in several species. The widely used jellyfish software appears to have no option but a very large -h (e.g -h 999999999999) to prevent data truncation. GSE-K software could and should check input histograms for truncation, and warn users of this mistake.

Contamination by non-nuclear DNA is a problem related to these low- and high-kmer contents, and can be addressed by excluding DNA that doesn’t map to a clean nuclear genome assembly, and/or by other means, e.g. Kraken (Wood et al 2019) software to classify taxonomic likelyhood of DNA reads, along with use of NCBI SRA’s taxonomic analyses (Katz et al 2021). These methods have been used with the results presented in this work.

In summary, GSE-Kmer methods are not measuring all of the genomic DNA, which can lead to under-estimates of genome sizes. Table DC8 compares the qualities of GSE-Gnodes and GSE-Kmer methods. Perfect-match tabulation of tiny sequences are at the core of these under-estimates. It may be difficult to correct the low-frequency ‘error’ problem, and reliably measure very-high copy repeats, without the imperfect matches possible with alignments of longer sequences.

GSE-Alignment methods as in Gnodes and others have an analogous ‘missing-genome’ problem: those reads un-aligned to an imperfect assembly of DNA, with map software that struggles to fully align duplicated DNA. Gnodes calculates GSE both ways: with and without un-aligned DNA, but prior and current results indicate that unaligned DNA is mostly part of the genome [Gnodes#2, #3, this]. Other GSE-A methods that exclude un-aligned DNA generally produce lower GSE. GSE-Kmer methods have a distinct advantage in speed and relative simplicity. Greater read depths and high sequence accuracy are needed for accurate kmer peak finding, whereas mapped reads can measure genome sizes, with declining precision, down to 1x coverage depth, using inaccurate long-reads as well as short. For rough-draft genome size estimates, GSE-K results are usually within about 10%, given good quality DNA of sufficient quantities (see Tables DC5,6 and related results).

For precision in assessing completeness of new genome assemblies, and in distinguishing genome changes among populations, the Gnodes type of measurement is more accurate. Gnodes agreement with the independent evidence of cytometric measures is a strong point in its favor.

For both GSE-A and GSE-K, artifactual biases in DNA shotgun sequences cause accuracy problems. Determining if such biases are large enough to affect measures is not simple, as artifact and genome biology often look alike. One reasonable solution is use of two or more DNA sequencing technologies and methods, and/or cytometric measures for comparison. Prior work [Gnodes #2] indicates that ONT long reads produce less-biased measures, in agreement with other evidence [please cite]. Illumina short reads have greater inconsistency, over a larger published set of data, than PacBio and ONT long reads; PacBio lofi versus hifi have different biases, the later hifi appears to have a bias against duplications due to computational averaging.

**Table DC8.**
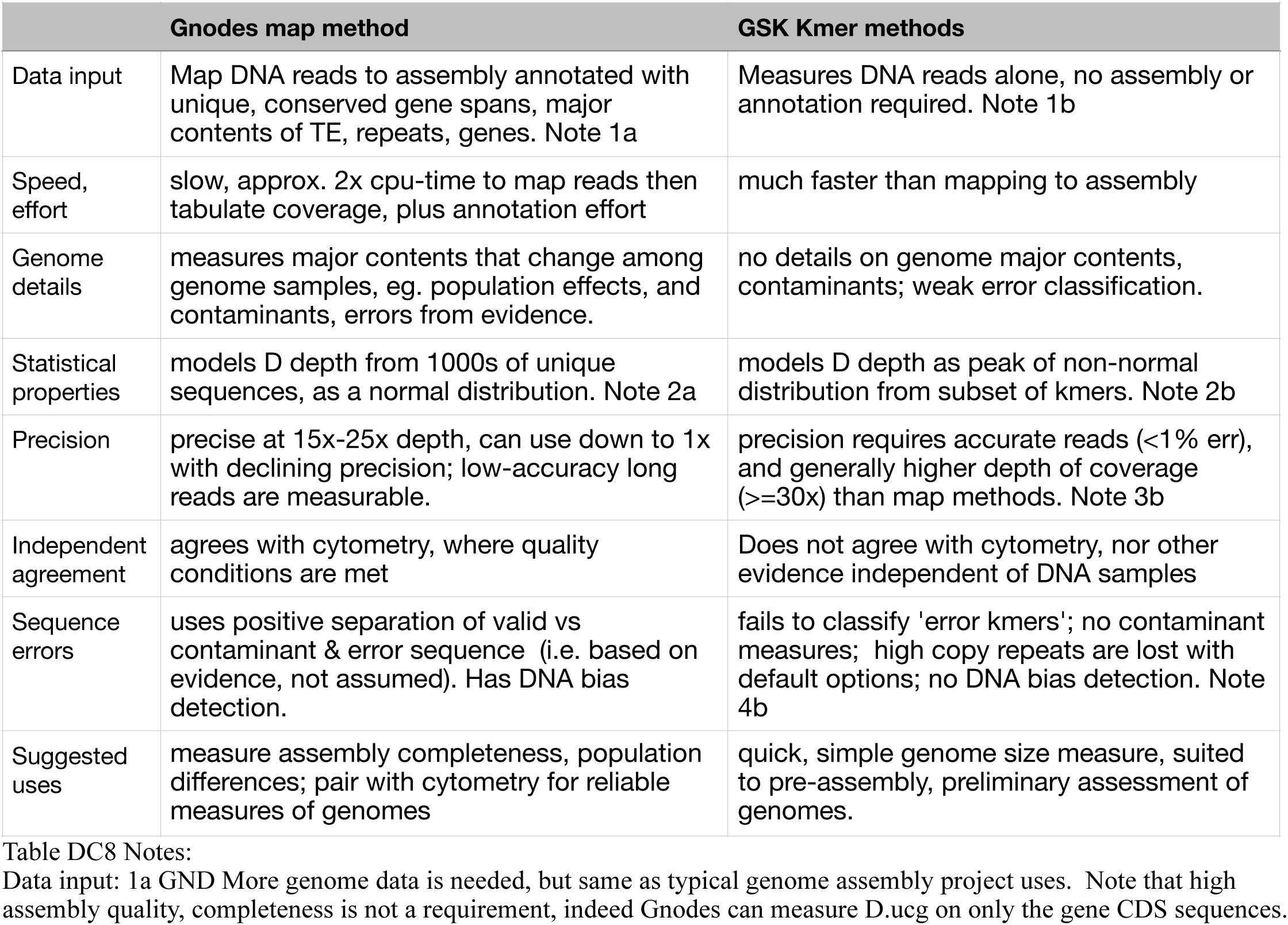
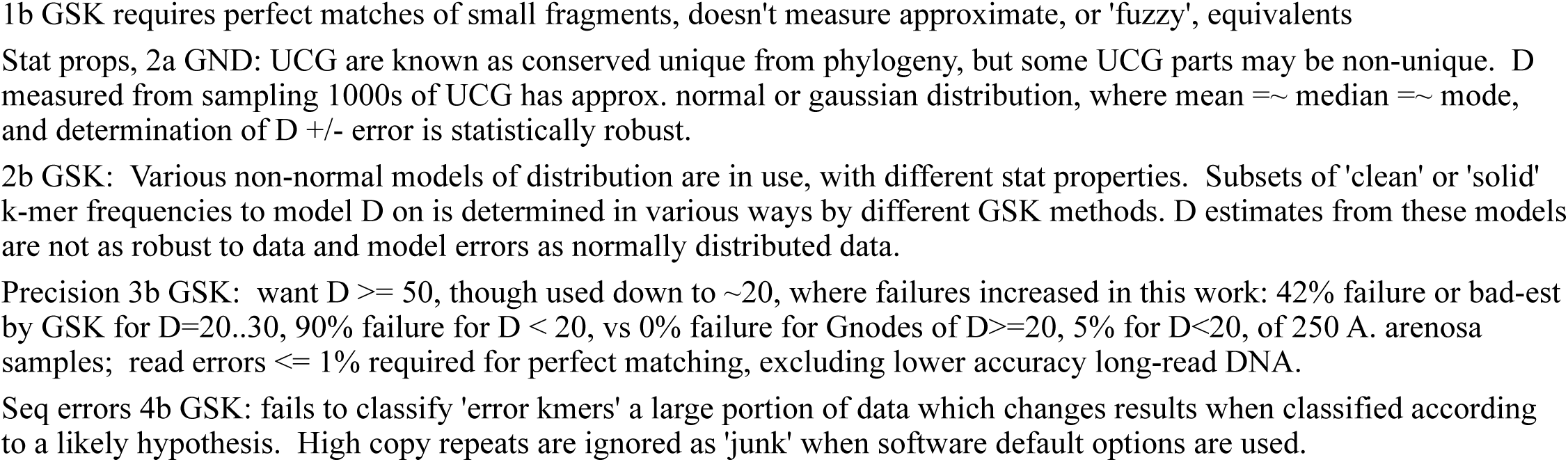
Genome measuring attributes of Gnodes map vs GSK kmer methods.

## Discussion

Uncertainty remains in genome size measurements, by cytometry, shotgun DNA reads, and other methods. However, comparisons of DNA reads and cytometry from available public data warrant a conclusion that these are finding the same genome sizes. These methods are precise enough, when care is taken to reduce errors, to detect population differences in genome sizes, consistent with regional and environmental clines (e.g Igolkina et al. 2025).

Much of the discrepancy in genome assemblies falling below cytometry is ascribed to duplicated or repetitive regions of genomes that remain difficult to assemble. It is possible to obtain assemblies matching the sizes measured by CYM and DNA methods, when multiple pieces of duplicated regions are retained.

Large duplicated genome regions fluctuate in size within species, not only among regional and inbred populations, but within individuals at different stages of life [human genome tandem repeat contents change with age, Ershova et al 2025], or different cell types [fruitfly thorax cells underreplicate repeat DNA, Hjelmen et al 2020]. Biological reasons and effects of this are being examined; accurate measures of such changes are of value.

Genome size measures with cytometry are subject to several methodological effects (e.g. Nix et al 2024), but no evidence of a systematic measurement bias for those has been established (Temsch et al. 2021). Determining the most accurate and reliable protocols for genome measurements with cytometry and DNA reads can be done by using both method to cross-validate their results.

Measurements on shotgun DNA need to take account of several error sources, including laboratory methods involving biosamples, DNA collection, DNA sequencing technology, as well as statistical methods for measuring this DNA. A review of sequencing and methodological problems with duplicated and repetitive DNA (Cechova 2021) has identified several sequencing technology factors that affect these measures, indicating that each method has its own problems and values, but in general they agree with each other, and are all “probably correct”. Prior and current results with Gnodes measures (Gnodes#2, this doc) are conclusive that several DNA processing biases reduce duplications. DNA per-base error correction methods are collapsing natural duplication variance into averaged sequences. DNA filters applied for presumed complications such as heterozygosity and artifacts are reducing or ignoring natural duplicated DNA.

Results here suggest the best measurement choices include Oxford Nanopore sequencing technology, for low bias in recovering duplicated DNA, and alignment methods to fully measure DNA coverage depth at the unit of unique genes and of duplicated regions. Biases in DNA technologies of Illumina short-reads, PacBio hifi and lofi complicate measurement. Genome content that is unaccounted for by kmer DNA perfect-match measures makes them less accurate.

### Recommendations for measuring genome sizes and contents

Use Ontl, as least biased long-read type, to measure genome sizes and contents most accurately.

Use two DNA types to estimate the range in measures.

Use cytometric measures, quantitative PCR and/or other types of genome size measures for comparisons.

Measure contents during steps in DNA filtering for assembly. Comparison of unfiltered versus filtered DNA, with assembly contents, is a quality check for completeness of genome assembly.

### Conclusions

Genome sizes from assemblies of animals and plants are often smaller than cytometric measures. Careful measures of DNA sequences generally support the larger cytometric measures. To resolve this problem of accurate genome measurement, new efforts are warranted to compare cytometric and DNA measures of suitable population-variable species with controlled laboratory techniques.

## Methods in detail

The major contents of genomes presented in these results are computed with Gnodes, as described in Gnodes#1 (Gilbert 2022). Gnodes software is part of the EvidentialGene package, publicly available at http://eugenes.org/EvidentialGene/ and http://sourceforge.net/projects/evidentialgene/ [Gilbert, 2019].

The basic algorithm of Gnodes is (a) align DNA reads to gene and chromosome assembly sequences, recording all multiple and unique map locations, (b) tabulate DNA cover depth at each sequence bin location for multiple and unique mappings, (c) measure statistical moments of coverage depth per item, itemized by categories of gene-cds, transposons and repeat annotations, duplicate and unique mappings, (d) summarize coverage and annotation tables, at chromosome-assembly and gene levels. These steps are summarized in the Box DC12 example. The statistical population of coverage is non-normal, so that median, average, skew and other statistics are calculated to approach precise and reliable measures. Table DC9 has moment statistic of three coverage depth measures for recent ONT R10 DNA samples.

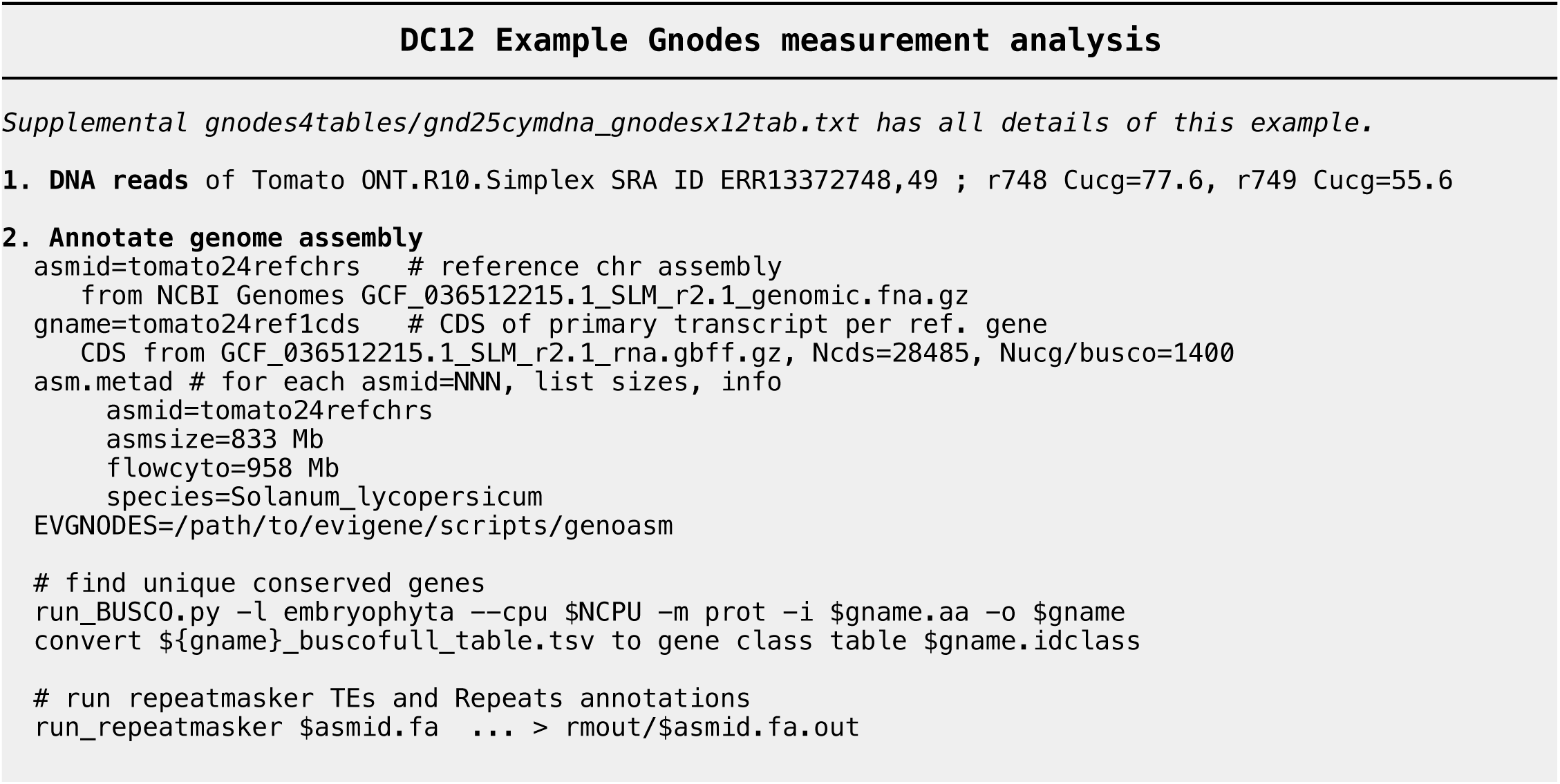
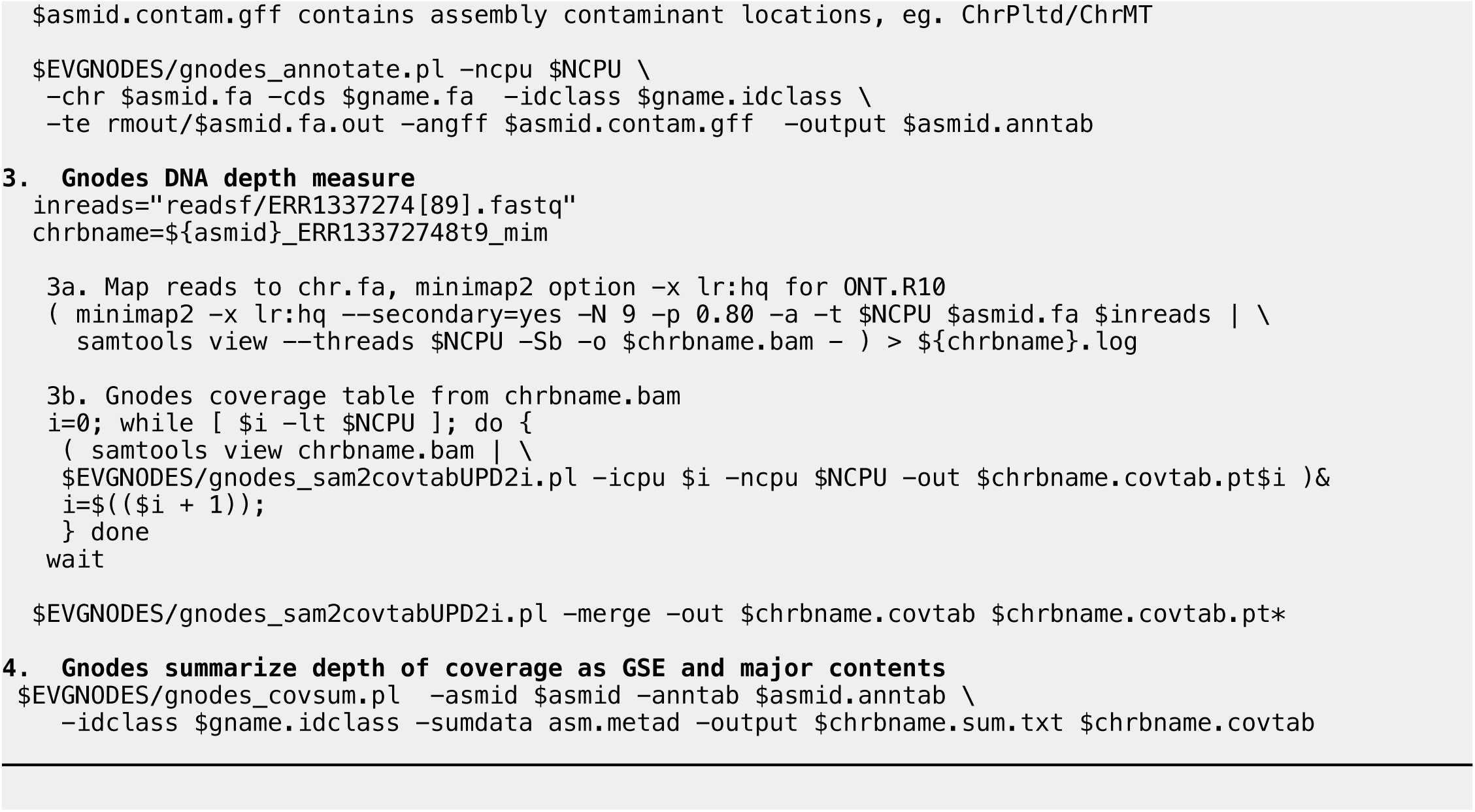

Updates to Gnodes software with this report include additions of depth of coverage density plots, as per Figures DC4 et seq.; a chromosome ploidy input option, for more accurate measures where some chromosomes, eg. haploid sex, have different ploidy levels; addition of a simplified depth of coverage tabulator, gnodes_sam2covupd2i.pl, better suited to interpreting the algorithm than the more complete gnodes_sam2covtab.pl. Table DC9 has statistics of depth estimates calculated with Gnodes for ONT R10 DNA samples as found in Table DC6. The UCGmed measure is approximately normally distributed, with average close to median, and little skew or kurtosis, depending on DNA qualities. These attributes do not ensure the UCGmed estimate is a true depth value, but depth extremes and unbalanced depths are reduced, and the close to normal distribution suggests a best estimator with random errors. However as noted in results above, when DNA samples are biased in some way, e.g. GC-bias or deficient in repeats, noted when C.all < C.ucg, then depth over all the assembly (D.est = allasm, or C.all) is a better estimator for the complete genome. In the putatively unbiased DNA samples of Table DC9, median C.all >= C.ucg, the expectation where C.all contains duplications, denoted also by large positive skew and kurtosis values.

**Table DC9.** Moment statistics of Gnodes depth estimates for ONT R10 Simplex DNA samples. UCGmed is median depth per unique gene, for n=300 to 2000 genes; CDSbus is depth over all annotated unique gene locations; D.est allasm is for all assembly locations, in 100b bins, asm size = 100*n. pnorm is p-value of Shapiro-Wilks test of non-normality.

| SppID | D.est | med | ave | sd | skew | kurt | n | pnorm | DNA_SRA_ID |
| --- | --- | --- | --- | --- | --- | --- | --- | --- | --- |
| A Arath | UCGmed | 89.5 | 88.6 | 13.2 | -0.36 | 0.37 | 298 | 0.04 | ERR7919757 |
| T Tomato | UCGmed | 56 | 55.4 | 9.84 | -0.05 | -0.002 | 1048 | 0.29 | ERR13372749 |
| M Maize | UCGmed | 55 | 55.7 | 8.35 | 0.20 | -0.30 | 505 | 0.02 | ERR13381137 |
| F Finch | UCGmed | 52 | 52.3 | 7.66 | 0.06 | 0.04 | 1701 | 0.01 | SRR32887368t9 |
| Z zFish | UCGmed | 45 | 45.4 | 6.55 | 0.16 | 0.46 | 2245 | 1e-07 | ERR12118113 |
| SppID | D.est | med | ave | sd | skew | kurt | n | pnorm | DNA_SRA_ID |
| A Arath | CDSbus | 93 | 92.9 | 12.3 | -0.13 | 0.57 | 29400 | 3e-08 | ERR7919757 |
| T Tomato | CDSbus | 56 | 56.2 | 9.68 | -0.05 | 0.12 | 30746 | 5e-04 | ERR13372749 |
| M Maize | CDSbus | 59 | 59.9 | 10.17 | 0.01 | 1.71 | 33217 | 1e-20 | ERR13381137 |
| F Finch | CDSbus | 53 | 53.3 | 8.38 | 0.17 | 0.52 | 63803 | 3e-05 | SRR32887368t9 |
| Z zFish | CDSbus | 46 | 45.9 | 7.93 | 0.08 | 0.46 | 66152 | 2e-08 | ERR12118113 |
| SppID | D.est | med | ave | sd | skew | kurt | n | pnorm | DNA_SRA_ID |
| A Arath | allasm | 89 | 87.7 | 17.4 | 0.10 | 6.17 | 1422k | 3e-30 | ERR7919757 |
| T Tomato | allasm | 58 | 57.9 | 15.50 | 2.62 | 24.73 | 8017k | 6e-35 | ERR13372749 |
| M Maize | allasm | 62 | 62.5 | 13.25 | 3.33 | 35.01 | 21688k | 1e-26 | ERR13381137 |
| F Finch | allasm | 52 | 52.3 | 11.70 | 2.46 | 32.26 | 11382k | 3e-17 | SRR32887368t9 |
| Z zFish | allasm | 45 | 44.9 | 11.09 | 1.63 | 25.07 | 14262k | 3e-25 | ERR12118113 |

### Data sets: DNA read sets, genome assemblies & annotations

DNA reads are drawn from public archives at NCBI SRA, and most chromosome assemblies from NCBI Genomes public archive. Supplemental tables for this paper, in gnodes4tables subfolder, include tables per species of GSE, with columns of Popnam, SRA_Reads, FCmb, Gnodes and other data, described in gnodes4tables/Gnodes4Tables_readme.txt

### Methods of DNA mapping

The author has examined several long-read DNA mapping programs: minimap2 [https://doi:10.1093/bioinformatics/bty191], widely used, is the default mapping software for Gnodes. For short-read mapping, bwa-mem2 [arXiv: 1303.3997] and minimap2 are used.

### Methods for genome annotations

Sequences of reference annotations for species were used by preference, where available in good quality, for coding genes, transposons, and repeats. Reference gene CDS were extracted from gene data provided by NCBI Genomes [ncbi-genomes/GCF_accession*rna.gbff]. The outputs of RepeatMasker, and RepeatModeler for some cases, were used [Smit et al, http://www.repeatmasker.org]. Computations done for this project include runs of tandem repeat finder (TRF, Benson 1999) and high order repeat finder (SRF, Zhang et al 2023) for some genome assemblies. Annotations used with the Maize NAM isoline assemblies were produced for each isoline, in preference to use of reference Maize assembly, in accordance with the large isoline differences in genome contents.

Non-nuclear DNA contents are annotated also (chloroplast, mitochondria and contaminants, mostly bacteria), and tabulated in result tables, with their amounts subtracted from GSE values. For some measures, non-nuclear DNA was removed first, using only DNA reads aligned to nuclear assemblies. Foreign contaminants are identified by NCBI SRA reports from STAT (Katz et al. 2021) and/or Kraken (Wood et al 2019) for taxonomic analysis of DNA sequences.

### Methods for major genome content measures

Genome assemblies and major contents sequences of CDS, transposons, and repeats were processed with Gnodes, which uses BLAST alignment of these to produce genome location tables congruent with the genome read-coverage tables also produced by Gnodes. Component scripts of Gnodes summarizes and plots DNA copy numbers relative to genome assembly (xCopy), and megabase measurements for the major contents.

### Methods for figures and tables

Data used to produce figures and document tables for species and population samples, including DNA Read public IDs, GSE for Cym, Gnodes, and GSK methods, along with accessory information on major genome contents, DNA depth values, contaminant counts, are provided in supplementary gnodes4tables/ folder, with descriptions in gnodes4tables/Gnodes4Tables_readme.txt. R scripts for most of the figures and statistics derived from these tables are found in gnodes4tables/gnd25cymdna_rstats.txt

### Methods for kmer error measurement

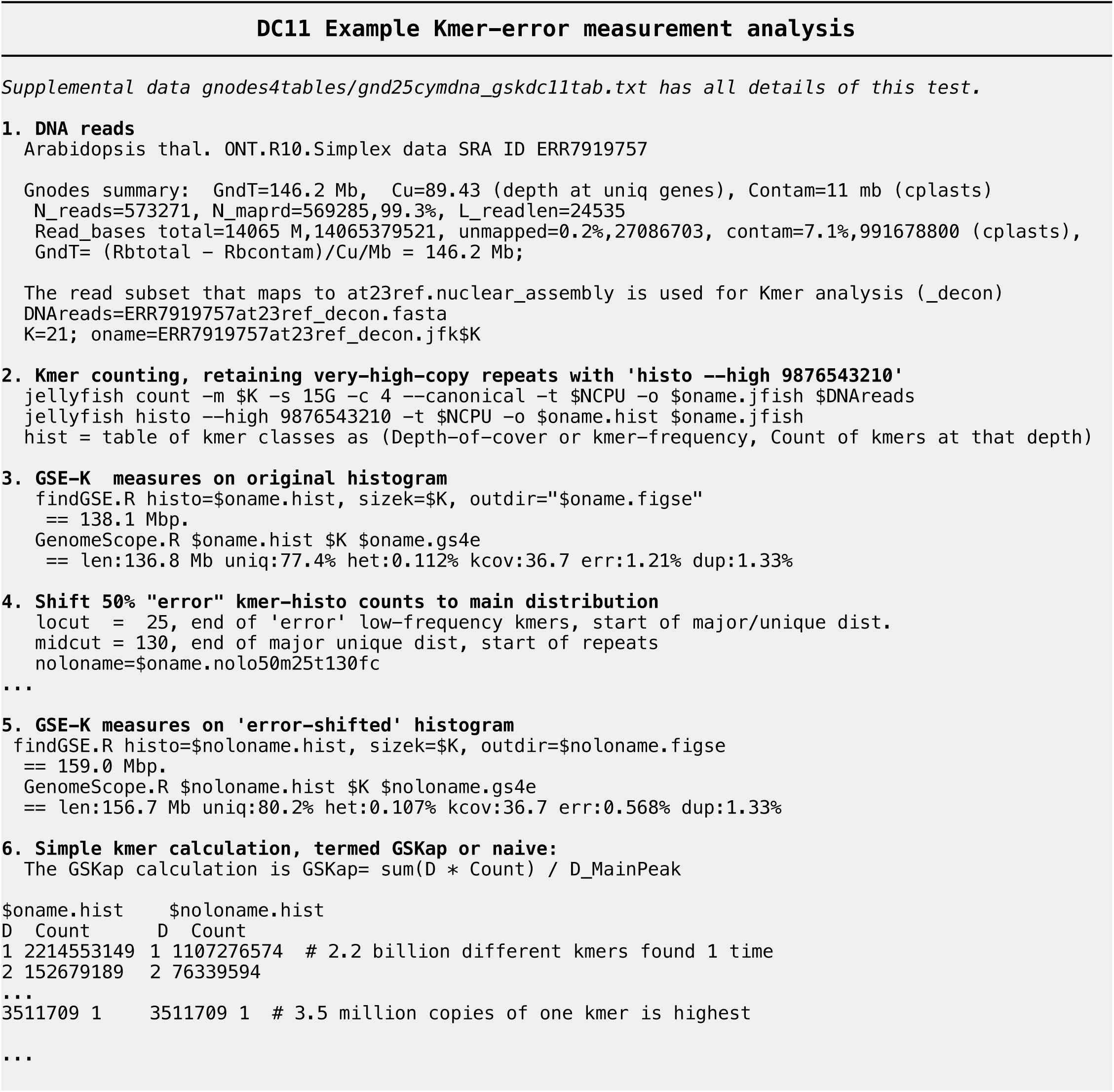
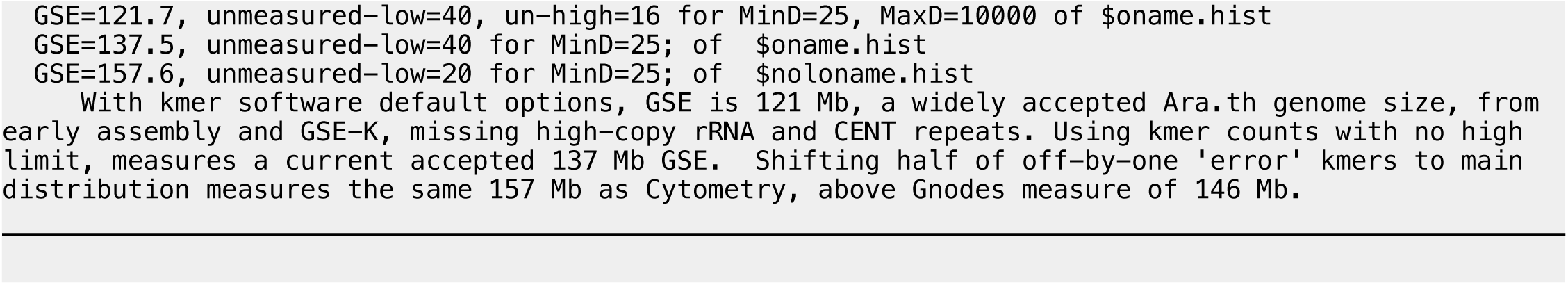

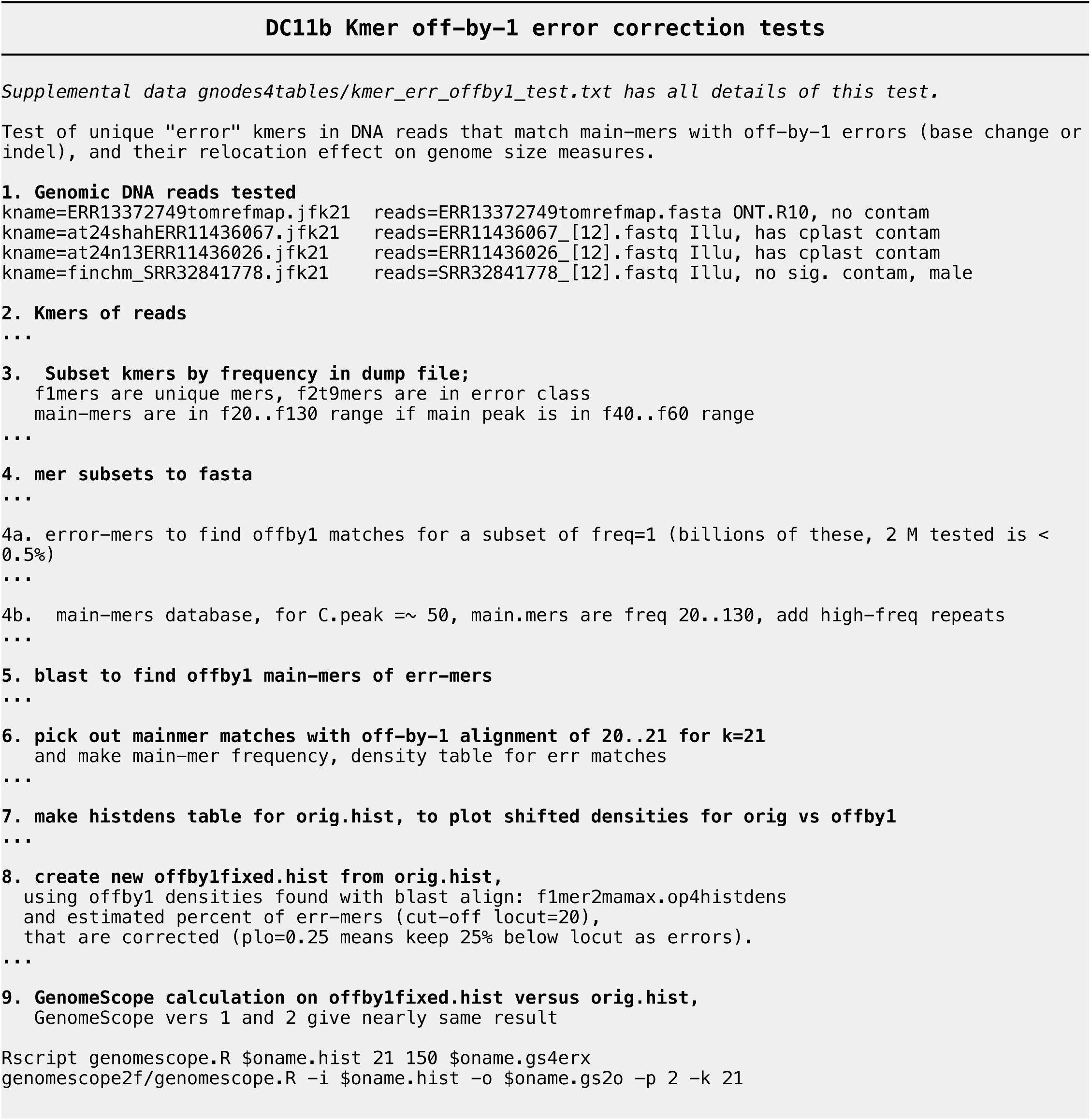
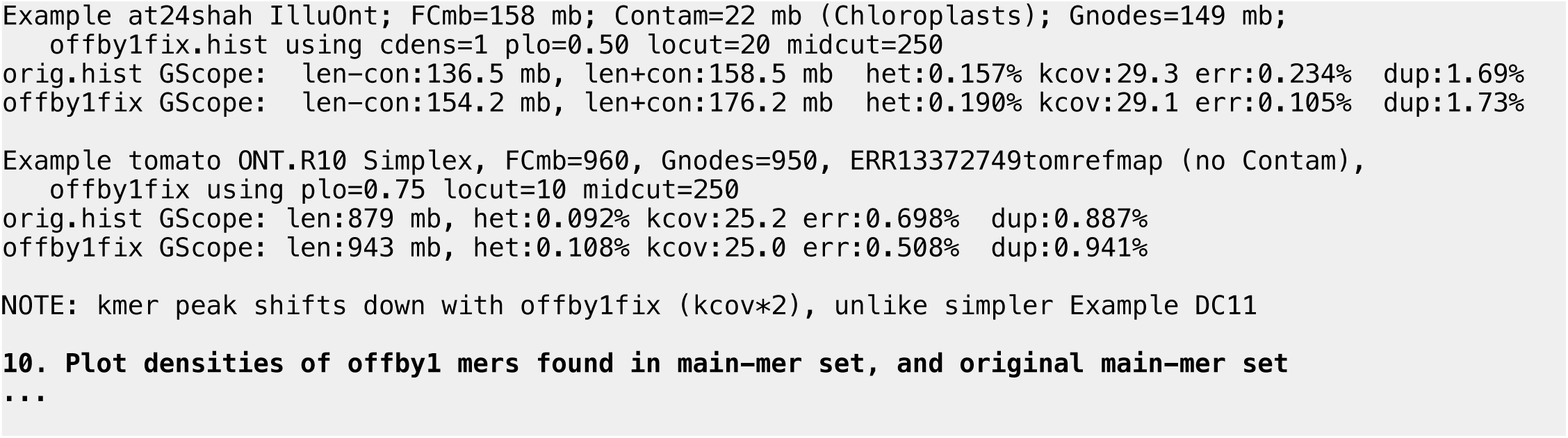

## Acknowledgements

My thanks to Filip Kolář who provided flow cytometry data of *Ara. arenosa* populations, as referenced in Kolar et al. 2015. The following provide shared computational resources to support this work: Advanced Cyberinfrastructure Coordination Ecosystem: Services & Support (ACCESS), funded by U.S. National Science Foundation, including resources at Indiana University (https://jetstream-cloud.org), Purdue University, and San Diego Supercomputer Center.

## Supplemental data

Data tables in directory gnodes4tables/ of http://eugenes.org/EvidentialGene/other/gnodes/gnodesdoc/

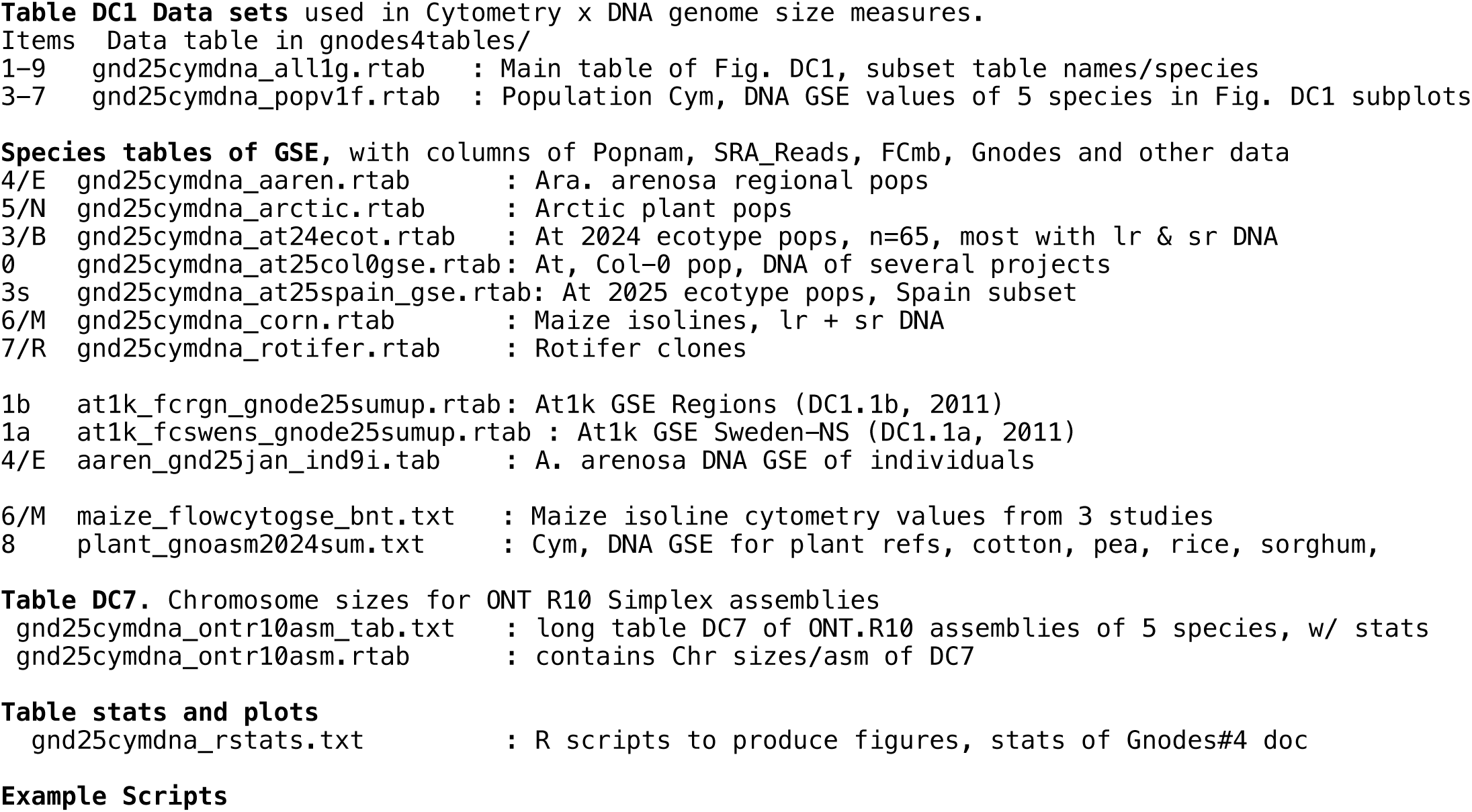
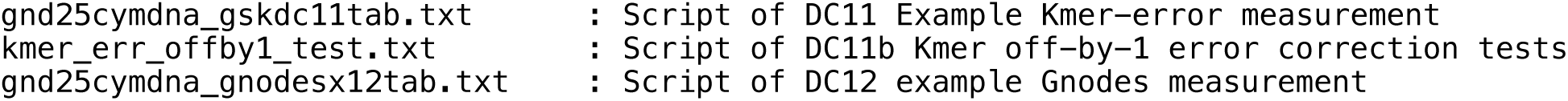

